# Differential TDP-43 interactomes between the cortex and cerebellum in the mouse

**DOI:** 10.64898/2026.01.16.699854

**Authors:** T Baldacchino, S Lynham, S Taconelli, C Vance, JC Mitchell

**Affiliations:** Department of Basic and Clinical Neuroscience, Institute of Psychiatry, Psychology and Neuroscience, King’s College London, Denmark Hill, London, SE5 8AF, UK; Centre of Excellence for Mass Spectrometry, The James Black Centre, King’s College London, London, UK

**Keywords:** TDP-43, mouse, cortex, cerebellum

## Abstract

TAR DNA binding protein 43 (TDP-43) is the core pathogenic protein across a spectrum of amyotrophic lateral sclerosis (ALS) and frontotemporal dementia (FTD) cases, but its pathological deposition shows regional selectivity, with abundant TDP-43 positive aggregates in the frontal cortex and spinal cord, and a much lower burden in the cerebellum. In health, TDP-43 expression in the cerebellum is higher than in cortex, hence what underpins this differential vulnerability to TDP-43 aggregation is unclear. Here we demonstrate that in healthy C57Bl/6J mice not only is TDP-43 expression higher in the cerebellum than the cortex, but that this expression difference is driven primarily by differences in cytoplasmic load. Mass spectrometry analysis of TDP-43 pull downs from the cortex and cerebellum of healthy mice identified TDP-43 interactors across a number of core functional pathways, with numerous differences between the two brain regions. Data are available via ProteomeXchange with identifier PXD062532. Notably, there were more interactors identified in both nuclear and cytoplasmic fractions within the cortex than the cerebellum. Putative interactions with four core paraspeckle proteins; SFPQ, NONO, FUS and PSPC1 were confirmed using immunoprecipitation and western blot analysis. Follow up validation using proximity ligation assay showed abundant perinuclear cytoplasmic interactions between TDP-43 and all four paraspeckle proteins in both large motor cortex neurons and purkinje cells, with significantly reduced nuclear interactions detected in the motor cortex for SFPQ, FUS and PSPC1. These findings suggest that TDP-43-protein interactions markedly differ between the TDP-43 pathogenesis vulnerable cortex and relatively resistant cerebellum and exploring these differences may yield new insight into disease mechanisms within ALS/FTD.

## Introduction

Aberrant cytoplasmic accumulation of TDP-43 (TAR DNA binding protein 43) is a core feature of around 97% of amyotrophic lateral sclerosis (ALS) and 50% of frontotemporal dementia (FTD) cases [1, 2]. Mutations in the *TARDBP* gene that codes for the protein have also been identified in a small subset of cases [3–5], while a number of disease causative mutations in other genes, including the most prevalent *C9orf72* expansion mutation, also present with TDP-43 aggregation as a core pathogenic feature [6–8], highlighting its integral role in the disease process.

As with a number of neurodegenerative disorders, ALS and FTD are characterised by regional variations in pathology load and neuronal loss, with the motor cortex and spinal cord primary targets in ALS, and the frontal and temporal lobes primary targets in FTD. In both cases, TDP-43 pathology is abundant in these affected regions [9, 10].

In contrast, despite evidence of significant crosstalk between the cortex and cerebellum [11–13], the pathogenic TDP-43 burden in the cerebellum is low and only evident in late-stage disease [14] raising questions as to what may drive this differential disease vulnerability. The C9orf72 expansion mutation in particular highlights this intriguing question. In patients with this mutation, mutation specific RNA foci and p62 labelled di-peptide repeat accumulations are abundant in the cerebellum, but TDP-43 aggregates are largely absent [15–17]. While these mutation specific pathologies are associated with some impairment of cerebellar function [18] and studies are increasingly demonstrating evidence of focal cerebellar atrophy in both ALS and FTD [19, 20], it has been demonstrated that frank neuronal loss in the cerebellum is not apparent in sporadic cases in the absence of C9orf72 or ataxin 2 mutations [21].

Conversely, in the cortex and spinal cord, where TDP-43 accumulation is present, significant neuronal cell loss is observed [22, 23], suggesting that TDP-43 pathology or associated consequences may drive cell death. The cerebellum has been reported to have markedly higher levels of TDP-43 than other regions of the cerebrum including the cortex [24] suggesting that reduced TDP-43 burden does not underpin this regional lack of pathology in disease.

TDP-43 is an RNA binding protein that shuttles between the nucleus and cytoplasm and is known to be involved in an array of functions, including RNA splicing [25–27] RNA transport, stress granule formation [28], and local protein synthesis in dendrites contributing to neuronal plasticity [29]. To date, a number of proteins have been proposed to interact with TDP-43, primarily falling into two families; nuclear proteins involved in RNA linked functions such as splicing and cytoplasmic proteins including translation initiation and elongation factors, and ribosomal subunits [30]. More recent studies have also identified TDP-43 interactions with several mitochondrial proteins [31] highlighting the complex functional profile of this protein.

A study exploring TDP-43 interactors in the human spinal cord found some unique proteins not identified in previous studies, suggesting that there may be tissue specific differences in TDP-43 interactors within the central nervous system [32]. It is possible therefore that tissue differences in the functional interactions of TDP-43 may underpin the differential vulnerability of cells within the central nervous system to TDP-43 pathology development.

In this study, we examine differences in TDP-43 expression and localisation between the cortex and cerebellum of healthy control mice, showing increases in protein expression within the cerebellum, with a relative increase in cytoplasmic load. We conduct mass spectrometry analysis to identify differences in the TDP-43 interactome between the two brain regions with follow-up validation experiments showing regional differences in the interactions of TDP-43 with a number of paraspeckle proteins within the cytoplasm of the cortex and cerebellum. We hypothesize that these distinct interaction profiles provide insight into the differential vulnerability of cortical and cerebellar cells to TDP-43 pathology.

## Methods

### Ethics Statement

All experiments were performed under the terms of the UK Animals (Scientific Procedures) Act 1986, and were approved by the Kings College, London ethics review panel.

### Animals

C57BL/6J mice were used for all experiments. Animals were bred in house or obtained from Charles River (UK) and maintained on standard mouse chow (PicoLab® Rodent Diet 20 5053) and water ad libitum. They were housed in a specific pathogen free facility in a temperature and humidity-controlled environment at 21 ± 2°C on a 12h light/dark cycle. As far as possible, animals were socially housed with same sex littermates in groups of 2-5 mice per cage and aged to 3, 6, 12, 18 and 24 months as required.

### Western Blotting

For detection of TDP-43 levels, four mice at each age point (2 male and 2 female) were culled by cervical dislocation. Cortex and cerebellar regions were carefully dissected out and immediately snap frozen and stored at −80°C. Samples were then processed into nuclear and cytoplasmic fractions. Briefly, samples were weighed, homogenised in 5x weight/volume of cell lysis buffer (10mM Hepes, 10mM NaCl, 1mM KH_2_PO_4_, 5mM NaHCO_3_, 5mM EDTA, 1mM CaCl_2_, 1mM MgCl_2_). A small volume of lysate was harvested to serve as a total lysate sample and the remainder was centrifuged at 6300xg at 4°C for 10 minutes. Supernatant was harvested as the cytoplasmic fraction and the pellet was washed four times by resuspending in 1ml Tris/sucrose/EDTA buffer (TSE: 10mM Tris, 300mM sucrose, 1mM EDTA, 0.1% IGEPAL) and centrifuged at 4000xg at 4°C for 5 minutes. The pellet was then resuspended in TSE as the nuclear fraction. Protein samples were separated using a 4-12% Polyacrylamide Gel (ThermoFisher, #WG1403A) and then transferred to a nitrocellulose membrane (ThermoFisher, #IB23001). TDP-43 protein levels were assessed using a rabbit antibody raised to the C-terminal of TDP-43 (1:1000, Proteintech; 12892-1-AP). Mouse anti-GAPDH (1:5000, Sigma; G8795) was used as total lysate and cytoplasm loading control, while mouse anti-Lamin B1 (1:1000, Proteintech; 66095-1-1g) was used as a nuclear loading control. Anti-mouse and anti-rabbit secondary antibodies conjugated to DyLight^TM^ 680 or DyLight^TM^ 800 (1:5000, Thermo-Fisher) were used to detect protein bands and visualised using the Odyssey Imager (Licor). For detection of paraspeckle protein levels, 3-month-old mice (n=3) were culled and cortex and cerebellum samples were harvested, fractionated and processed as described above. Mouse anti FUS (1:500, Santa Cruz: sc-373698), rabbit anti-PSPC1 (1:1000, Abcam: ab133574), mouse anti-SFPQ (1:500, Invitrogen: MA1-25325) and rabbit anti-NONO (1:500, Abcam: ab133574) antibodies were used with loading controls and secondary antibodies as before.

### Immunohistochemistry

For assessment of TDP-43 localisation and expression, young (3-month, n=3) and aged (24-month, n=3) mice were deeply anaesthetised and transcardially perfused with ice cold PBS followed by 4% paraformaldehyde. Brains were dissected out and postfixed in 4 % PFA in 15% sucrose for 16h, cryoprotected in 30 % sucrose for 24 h and cut into 30μm coronal sections on a cryostat. For immunohistochemistry, the following antibodies were used: Rabbit anti-TDP (C-terminal; 1:400, Proteintech 12892-1-AP), rat anti-TDP (1:400, Biolegend SIG-39854; for co-staining with paraspeckle proteins), chicken anti-MAP2 (1:1000, Abcam, #ab9241489), rabbit anti-FUS (1:200, Bethyl A300-294A), rabbit anti-SFPQ (1:400, Abcam ab177149), rabbit anti-PSPC1 (1:200, Santa Cruz sc-374181), rabbit anti-NONO (1:250, Abcam ab133574). Following overnight incubation at 4°C in the relevant primary antibodies, free-floating sections were washed and incubated with secondary antibodies (donkey anti-rabbit AlexaFluor^TM^ 488, donkey anti-rat AlexaFluor^TM^ 568 (where required), goat anti-chicken AlexaFluor^TM^ 647). Sections were then washed and incubated in DAPI (4ʹ, 6-Diamidino-2-phenylindole dihydrochloride, ThermoFisher, #62248) for 10 minutes, prior to a final wash, and subsequent mounting onto glass slides using FluorSave (Merck, #345789).

For TDP-43 intensity assessment, motor cortex and cerebellum (Crus1 and paraflocculus) were imaged using an Opera Phenix^TM^ screening system (Perkin Elmer) with subsequent analysis using the Harmony^TM^ imaging and analysis software. For TDP-43 and paraspeckle colocalization studies, samples were imaged using a Nikon iSIM super resolution microscope.

### Immunoprecipitation

Cortex and cerebellum were harvested from 3-month old C57Bl/6J mice (4 male, 4 female). To allow sufficient tissue for fractionation and subsequent processing, samples were pooled such that each n number contained one male and one female mouse. Samples were fractionated as described above then incubated with rabbit polyclonal N-terminal antibody (Abcam #ab225710) or rabbit polyclonal C-terminal antibody (Proteintech #12892-1-AP) coupled to Protein G Dynabeads (ThermoFisher, #10003D) at 4°C overnight (5μg/ml for nuclear fraction and 20μg/ml for cytoplasmic fraction). Following validation of successful pull-down via western blot detection of TDP-43 as described above, samples were prepared for mass spectrometry analysis. Samples were run on a NuPAGE™ 4-12% Bis-Tris Midi Protein Gel (ThermoFisher, #WG1403BOX) for ∼10min, stained with Coomassie (Abcam, #ab119211) and sent to the KCL Proteomics facility for mass spectrometry analysis.

For co-immunoprecipitation studies, immunoprecipitation and flow-through samples were separated using NuPAGE™ 4-12% Bis-Tris Midi Protein Gels and transferred onto a nitrocellulose membrane. Western blot detection was conducted as described above. Primary antibodies used were: mouse anti-FUS (1:500; Santa Cruz sc-373698), rabbit anti-PSPC1 (1:1000, Abcam ab133574), mouse anti-SFPQ (1:500, Invitrogen MA1-25325) and rabbit anti-NONO (1:1000, Abcam ab133574).

### Mass Spectrometry

Coomassie stained gel sections were destained using 50% acetonitrile (ACN):50% 100mM triethylammonium bicarbonate (TEAB) prior to enzymatic digestion. Proteins were reduced with 10mM dithiothreitol (DTT; D9779, Sigma) at 56°C followed by alkylation with 55mM iodoacetamide (IAM; I1149, Sigma) at room temperature in the dark to form stable carbamidomethyl derivatives at cysteine residues. Proteins were enzymatically digested using trypsin in an enzyme:substrate ratio of 1:80. Peptides were extracted from the gel pieces and dried to completion in a SpeedVac (Thermo Fisher Scientific).

Extracted peptides were resuspended in 2% ACN/0.05% trifluoroacetic acid prior to analysis by high-resolution Orbitrap mass spectrometry coupled to nano-UHPLC. Chromatographic separation was performed using a U3000 UHPLC NanoLC system (ThermoFisherScientific, UK). Peptides were resolved by reversed phase chromatography on a 75μm C18 Pepmap column (50cm length) using a three-step linear gradient of 80% acetonitrile in 0.1% formic acid. The gradient was delivered to elute the peptides at a flow rate of 250nl/min over 60 min starting at 5% B (0-5 minutes) and increasing solvent to 40% B (5-40 minutes) prior to a wash step at 99% B (40-45 minutes) followed by an equilibration step at 5% B (45-60 minutes).

The eluate was ionised by electrospray ionisation using an Orbitrap Fusion Lumos (ThermoFisherScientific, UK) operating under Xcalibur v4.3. The instrument was first programmed to acquire using an Orbitrap-Ion Trap method by defining a 3s cycle time between a full MS scan and MS/MS fragmentation by collision induced dissociation. Orbitrap spectra (FTMS1) were collected at a resolution of 120,000 over a scan range of m/z 375-1600 with an automatic gain control (AGC) setting of 4.0e5 (100%) with a maximum injection time of 35 ms. Monoisotopic precursor ions were filtered using charge state (+2 to +7) with an intensity threshold set between 5.0e3 to 1.0e20 and a dynamic exclusion window of 35s ± 10 ppm. MS2 precursor ions were isolated in the quadrupole set to a mass width filter of 1.6 m/z. Ion trap fragmentation spectra (ITMS2) were collected with an AGC target setting of 1.0e4 (100%) with a maximum injection time of 35 ms with CID collision energy set at 35%.

### Database Searching

Raw mass spectrometry data were processed into peak list files using Proteome Discoverer (ThermoScientific; v2.3). The raw data file was processed and searched using the Mascot (www.matrixscience.com) & Sequest [33] search algorithms against the Uniprot Mouse Taxonomy (17,082 entries) database. All database searching was performed at a stringency of 1% FDR including a decoy database search. The database output file was uploaded into Scaffold software (version 5.0.1; www.proteomesoftware.com) for visualisation and manual verification.

### Proximity Ligation Assay

In situ protein-protein interaction assessment in mouse cortex and cerebellar samples was performed using the Duolink® Proximity Ligation Assay (PLA). An anti-rat antibody (Jackson’s ImmunoResearch, #712-005-153) specific MINUS probe was generated using the Duolink® In Situ Probemaker MINUS (Merck, #DUO92010), to allow avoidance of potential cross-reactivity arising from the use of a mouse-based probe and used in conjunction with the commercially available Duolink® rabbit PLUS probe (Merck #DUO92002). Following transcardial perfusion and fixation, brains were harvested from 3 C57Bl/6J mice and sectioned into 30μm thick sections using a cryostat. Free-floating sections containing the motor cortex and cerebellum were permeabilised in 0.1% Triton X-100 for 10 minutes, washed in sterile PBS and incubated in Duolink® blocking solution (Merck #DUO8200) for 1 hour at 37°C. Sections were then incubated overnight at 4°C in rat anti-TDP-43 (1:400, Biolegend #SIG-39854) together with a rabbit antibody to one of the paraspeckle proteins (FUS: 1:200, Bethyl A300-294A; PSPC1: 1:200, Abcam ab133574; SFPQ: 1:400, Abcam ab177149; NONO: 1:250, Abcam ab133574), in Duolink® antibody diluent (Merck #DUO82008). Samples were then washed three times for 10 minutes each time in Wash A buffer (Merck, #DUO82049) at room temperature and incubated in the rabbit PLUS and rat MINUS probes (1:5 and 1:20 respectively) in antibody diluent as per primary antibodies for 1 hour at 37°C. After a further three 10 minute washes in Wash A buffer, sections were incubated in ligation solution (Merck, #DUO92007) for 30 minutes at 37^0^C, then washed twice more in Wash A buffer. The rolling amplification hybridisation mix (Merck, #DUO92007) was then added and samples were incubated for 100 minutes at 37°C, with two final 10 minute washes in 1x Wash B buffer (Merck, #DUO82049) and one final 10 minute wash in 0.1x wash B buffer. Following this, samples were incubated in chicken anti-MAP2 (1:1000, Abcam, #ab9241489) at 4°C overnight, and underwent immunohistochemical staining using an anti-chicken 488 secondary antibody (Invitrogen, #A11059) as described above. Samples were imaged using a Nikon iSiM super resolution microscope. Four images of each region (layer V motor cortex, Purkinje cell layer, Granule cell layer) were taken for each sample and analysis was performed using Nikon NIS Elements.

### Statistical Analysis

For western blot analysis, quantification of bands was carried out using the Image Studio Lite application (Licor) using a median intensity and signal was normalised to the loading control. For TDP-43 expression levels, the resultant values were normalised against the mean data for the 3-month cortical sample. Data was then analysed statistically using GraphPad by way of two-way ANOVA followed by the post-hoc Tukey test, with age and brain region as the control variables. For paraspeckle protein analysis, data was analysed using GraphPad by way of Welch’s t-test.

For immunohistochemical analysis of TDP-43 expression in different cell populations, following initial imaging at 5x to identify the focal areas for analysis, focused images were taken using a 40x water lens. To identify neuronal cells, the inbuilt Harmony^TM^ AI training mode was used to train the system to recognise large and small neurons. Initially, nuclei were identified based on DAPI staining, then using the 647 channel, which detected MAP2, the software was trained to recognised patterns of shapes. Five defined populations emerged: large neurons, small neurons, dendrites, ‘large debris’ and ‘small debris’ (Figure 1f). Regions identified as ‘debris’ were not included in the analysis. Following training, samples were assessed for nuclear and cytoplasmic TDP-43 intensity in large and small cells, as well as total intensity in the ‘dendrites population. Resultant values were then analysed statistically using GraphPad by way of three-way ANOVA followed by the post-hoc Tukey test, with cell type, cell region (nuclear vs cytoplasmic) and age as the controlled variables.

**Fig 1.**
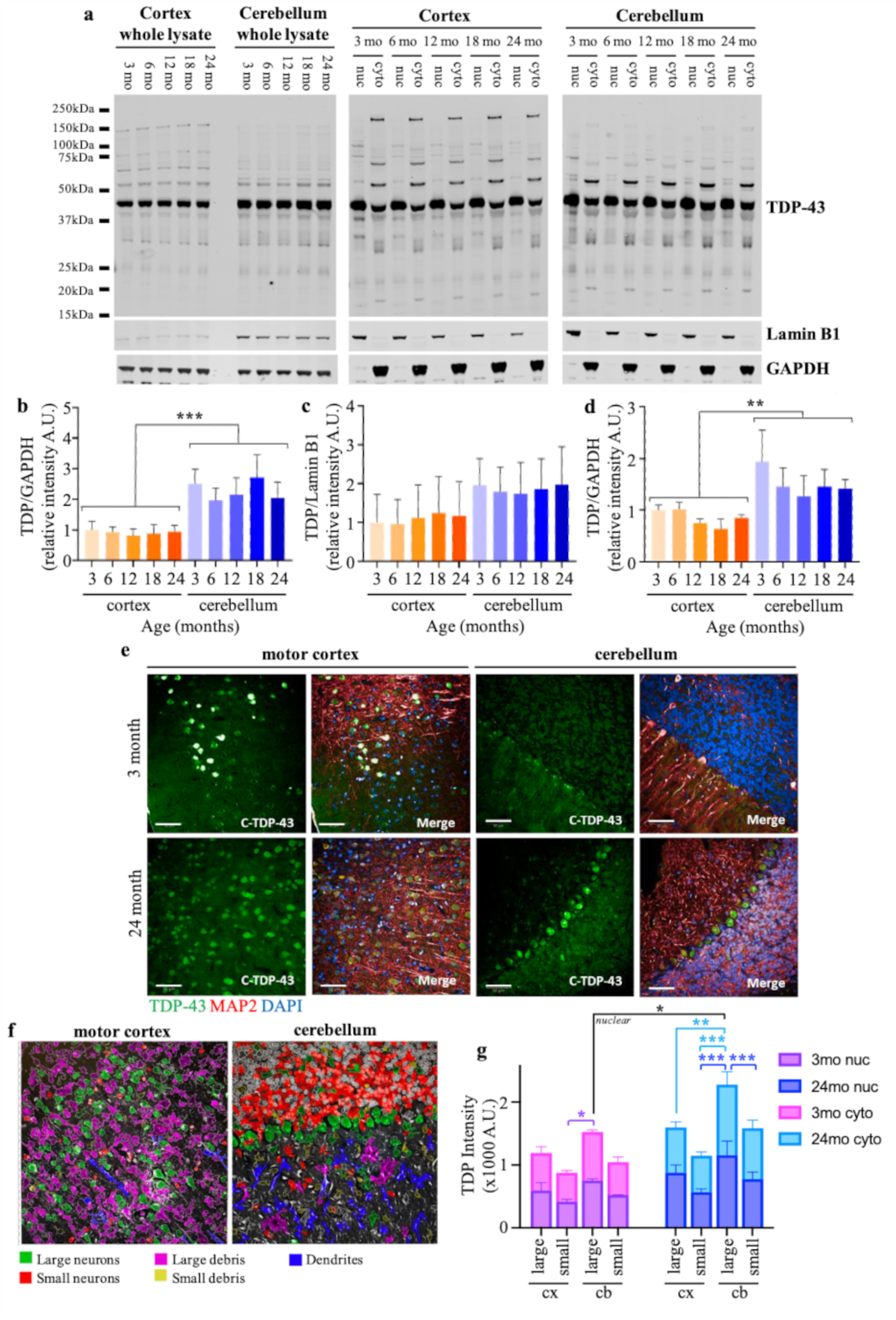
Cytoplasmic TDP-43 expression is enhanced in the cerebellum compared to the cortex. (a) Representative example of western blot assessment of nuclear and cytoplasmic TDP-43 levels within the cortex and cerebellum of C57Bl/6J mice throughout aging. (b-d) Quantitative analysis of TDP-43 expression from western blot analysis demonstrates a significant increase in total TDP levels in the cerebellum throughout aging (b), with no significant change specifically in the nuclear fraction (c), but a significant increase observed within the cytosolic fraction (d). (e) Representative examples of immunohistochemical detection of TDP-43 in the motor cortex and Crus 1 region of the cerebellum in young (3mo) and aged (24mo) C57Bl/6J mice. TDP-43 staining is shown in green. Neurons were detected using a MAP2 marker shown in red, and nuclei detected with DAPI are shown in blue. Scale bar is 50μm. (f) Representation of Harmony™ high-content analysis software detection of large and small neurons within the motor cortex and Crus1 region of the cerebellum following the inbuilt AI training pipeline focusing on MAP2 staining. (g) Quantification of TDP-43 intensities within the nucleus and cytoplasm of large and small neurons of the motor cortex and Crus1 cerebellum in young and aged C57Bl/6J mice. *p<0.05; **p<0.01; ***p<0.0001, 3-way ANOVA with Tukey post hoc

For PLA analysis, nucleus and cytoplasm binaries were established based on the presence or absence of DAPI. PLA signal was determined as any signal in the 568 channel with an intensity higher than the background. Mean PLA spot counts in the nuclear and cytoplasmic regions of each image was calculated, and mean spot size in each brain region was measured. Spot counts were analysed statistically using GraphPad by way of two-way ANOVA followed by the Fisher LSD post hoc test with brain region (motor cortex, Purkinje cell layer, granule cell layer) and cell region (nucleus, cytoplasm) as the controlled variables. Mean spot size in the different brain regions was analysed using one-way ANOVA with the Tukey post hoc test.

## Results

### TDP-43 expression is increased in cerebellar cytoplasm

Western blot analysis of TDP-43 within the cortex and cerebellum of non-transgenic C57Bl6/J mice demonstrated increased levels of the protein in the cerebellum regardless of mouse age (p<0.001; Fig 1a-b), consistent with previous reports [24]. This increase appeared to be primarily driven by an increase in cytoplasmic levels (p=0.0015; Fig 1d), while nuclear levels of TDP-43 were not significantly different between the two brain regions (Fig 1c; p=0.1427). Aging *per se* did not impact on either total TDP-43 levels, or nuclear-cytoplasmic localisation.

Immunohistochemical analysis of young (3 month) and aged (24 month) mouse motor cortex and cerebellum (Crus 1 and paraflocculus) displayed similar region-specific differences in both nuclear and cytoplasmic levels of TDP-43 (Fig 1e, g). Following Harmony™ high-content analysis software AI identification of large and small neurons in each region, as demonstrated in Fig 1f, three-way ANOVA assessment of age, cell type and cell region (nuclear vs cytoplasmic) demonstrated a significant impact of age (p<0.0001) and cell type (p<0.0001) on TDP-43 levels, although no significant interaction between these factors was observed (p=0.1086). Data suggested that both nuclear and cytoplasmic TDP-43 levels were generally highest in the large cells of the cerebellum (likely to be Purkinje cells). Most notably, cytoplasmic load in these cells in aged mice was significantly higher than that observed in the large cortical neurons (p=0.0056). Unlike the western blot analysis, age did appear to increase TDP-43 load, most notably in the nucleus of the large cerebellar cells (p=0.0314), with a similar, but non-significant trend in the cytoplasm (p=0.0791). It is likely that the difference between the western blot and immunofluorescent assessments is due to the increased focus on specific cell populations in the immunofluorescent analysis.

It is also important to note that while TDP-43 is frequently described as a predominantly nuclear protein, both western blot and immunofluorescent analysis clearly demonstrate robust protein expression within both the nucleus and the cytoplasm of all cells assessed, and the immunofluorescent analysis detected no overall significant difference in protein load between these two subcellular regions, suggesting that levels of cytoplasmic TDP-43 may be somewhat comparable to that of the nucleus.

### TDP-43 interactors vary between the cortex and cerebellum

TDP-43 immunoprecipitation was conducted on nuclear and cytoplasmic fractions prepared from cortex and cerebellum samples harvested from 3-month-old healthy control C57Bl6/J mice (male and female samples pooled; n=4). Independent immunoprecipitations were conducted using both N-(Abcam) and C-(Proteintech) terminal TDP-43 antibodies. Western blot analysis demonstrated robust pull down of TDP-43 in the nuclear fraction of both the cortex and cerebellum with both antibodies. As might be expected, a reduced level of TDP-43 was evident in the cytoplasmic pull downs, although bands were consistently detected with the C-terminal antibody (Figure 2a).

**Fig 2.**
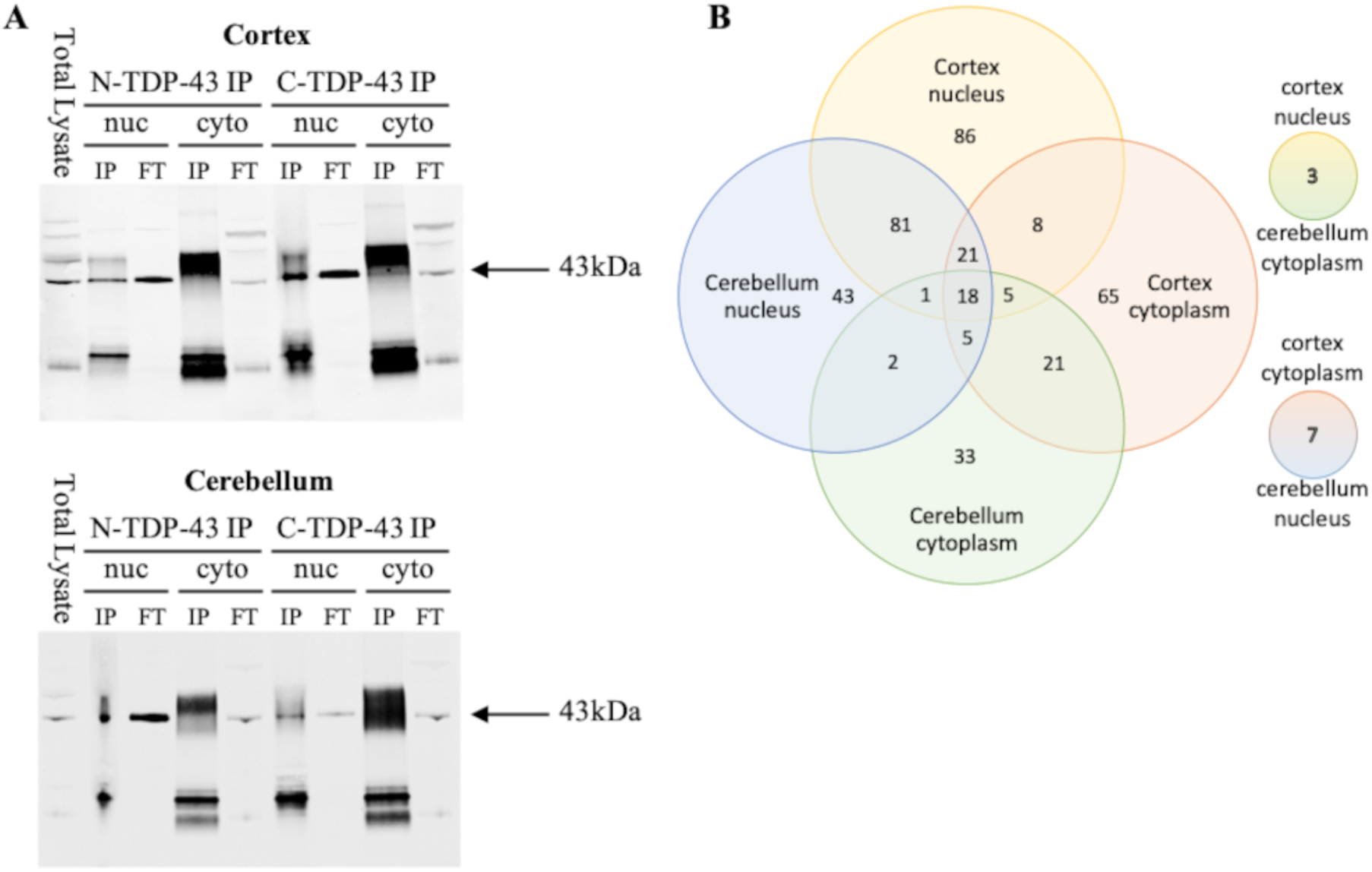
TDP-43 interactors differ between the cortex and cerebellum. (a) Antibodies raised to both the N (Abcam, #ab225710) and C (Proteintech #12892-1-AP) terminal of TDP-43 successfully pulled down TDP-43 from the nuclear fraction of both cortex and cerebellum samples. The C terminal antibody also successfully pulled down TDP-43 from the cytoplasmic fraction, albeit to a lesser extent. In contrast, no clear band was visible in cytoplasmic pull-downs using the N terminal antibody. IP: immunoprecipitation, FT: flow through. (b) Number of TDP-43 interactors identified, and their distribution across the various brain regions and cell fractions. While there were some overlaps between the cortex and cerebellum, there was marked differences in both nuclear and cytoplasmic interactors

Mass spectrometry analysis of these TDP-43 pull-downs identified ∼80-270 TDP-43 protein interactors per sample. The mass spectrometry proteomics data have been deposited to the ProteomeXchange Consortium via the PRIDE [34] partner repository with the dataset identifier PXD062532 and 10.6019/PXD062532. TDP-43 epitopes were detected in most samples, aside from a subset of N-terminal pull downs (Supplementary Table 1) which may indicate a poor quality pull down, or excessive fragmentation of the TDP-43 peptides during the digestion process.

The full list of TDP-43 protein interactors was filtered to only include those with 2 or more epitopes identified in a minimum of 2 samples. Following this, data from the two antibodies was pooled, and any apparent duplicates were removed, leaving a total of 399 different protein interactors for further analysis (Supplementary Figure 2). Of these, 321 were identified within the cortex, while only 240 were found in the cerebellum. Of the 321 proteins identified in cortex samples, 159 were not found in the cerebellum, while 78 of the 240 cerebellar interactors were not present in the cortex. This suggests there are core differences in the TDP-43 interactome between these two brain regions. As might be expected, given the known functions of TDP-43, a similar enrichment in interactors was identified in the nucleus (280 proteins total, 210 unique to nucleus), compared to the cytoplasm (189 proteins total, 119 unique to cytoplasm). A breakdown of the relative distribution of proteins within the different fractions assessed is provided in figure 2b.

STRING analysis [35] with subsequent MCL cluster analysis (inflation parameter=1.5) was carried out on the four data sets obtained. Six clusters were identified in the cortical nuclear samples (Supplementary Figure 3). In contrast, seven clusters were identified in cerebellar nuclear samples (Supplementary Figure 4), although one of these had no gene ontology biological processes associated with it. As might be expected, gene ontology assessment of the enriched biological processes found marked overlaps between the two datasets, with clusters associated with RNA processing, cytoskeletal organisation, chromatin organisation, and the extracellular matrix (Tables 1-2). However, there were some differences, with a small number of vesicular processes only enriched in the cortex, and small GTPase binding only enriched in the cerebellum.

**Table 1:** Top biological processes associated with each TDP-43 protein interactor in the cortex nuclear fraction detected using mass spectrometry analysis. Six clusters were identified using STRING analysis. Where more processes were enriched, the top 6 most significant processes are shown.

| Cluster | #term ID | term description | observed gene count | background gene count | strength | false discovery rate |
| --- | --- | --- | --- | --- | --- | --- |
| 1 | GO:0008380 | RNA splicing | 48 | 364 | 1.55 | 4.3E-58 |
|  | GO:0016071 | mRNA metabolic process | 53 | 613 | 1.37 | 7.77E-57 |
|  | GO:0006397 | mRNA processing | 48 | 452 | 1.46 | 2.29E-54 |
|  | GO:0006396 | RNA processing | 51 | 828 | 1.22 | 2.49E-47 |
|  | GO:0016070 | RNA metabolic process | 57 | 1498 | 1.01 | 1.65E-43 |
|  | GO:0000398 | mRNA splicing, via spliceosome | 35 | 231 | 1.61 | 1.45E-42 |
| 2 | GO:0007010 | Cytoskeleton organization | 43 | 1243 | 1.08 | 3.97E-34 |
|  | GO:0016043 | Cellular component organization | 55 | 5482 | 0.54 | 3.71E-21 |
|  | GO:0006996 | Organelle organization | 47 | 3482 | 0.67 | 5.4E-21 |
|  | GO:0000226 | Microtubule cytoskeleton organization | 25 | 559 | 1.19 | 8.05E-20 |
|  | GO:0007017 | Microtubule-based process | 27 | 840 | 1.05 | 3.05E-18 |
|  | GO:0030030 | Cell projection organization | 28 | 1259 | 0.89 | 4.99E-15 |
| 3 | GO:0051276 | Chromosome organization | 26 | 962 | 1.17 | 1.54E-21 |
|  | GO:0006325 | Chromatin organization | 22 | 599 | 1.3 | 3.58E-20 |
|  | GO:0006338 | Chromatin remodeling | 17 | 284 | 1.51 | 5.07E-18 |
|  | GO:0031497 | Chromatin assembly | 14 | 163 | 1.67 | 2.34E-16 |
|  | GO:0006996 | Organelle organization | 32 | 3482 | 0.7 | 2.17E-15 |
|  | GO:0034728 | Nucleosome organization | 13 | 143 | 1.7 | 2.17E-15 |
| 4 | GO:0110011 | Regulation of basement membrane organization | 5 | 11 | 2.82 | 4.99E-09 |
|  | GO:0007155 | Cell adhesion | 11 | 918 | 1.24 | 7.15E-09 |
|  | GO:2001046 | Positive regulation of integrin-mediated signaling pathway | 5 | 15 | 2.69 | 7.15E-09 |
|  | GO:0030198 | Extracellular matrix organization | 8 | 281 | 1.62 | 0.00000002 |
|  | GO:0051149 | Positive regulation of muscle cell differentiation | 5 | 80 | 1.96 | 0.00000405 |
|  | GO:0031589 | Cell-substrate adhesion | 6 | 197 | 1.65 | 0.00000441 |
| 5 | GO:0051301 | Cell division | 4 | 518 | 1.53 | 0.0185 |
|  | GO:0061640 | Cytoskeleton-dependent cytokinesis | 3 | 105 | 2.1 | 0.0185 |
|  | GO:1902855 | Regulation of non-motile cilium assembly | 2 | 17 | 2.71 | 0.0284 |
|  | GO:0022402 | Cell cycle process | 4 | 824 | 1.33 | 0.0315 |
| 6 | GO:0072583 | Clathrin-dependent endocytosis | 3 | 31 | 2.72 | 0.00022 |
|  | GO:0098884 | Postsynaptic neurotransmitter receptor internalization | 2 | 9 | 3.08 | 0.01 |
|  | GO:0099003 | Vesicle-mediated transport in synapse | 3 | 147 | 2.05 | 0.01 |

**Table 2:** Top biological processes associated with each TDP-43 protein interactor in the cerebellum nuclear fraction detected using mass spectrometry analysis. Seven clusters were identified using STRING analysis, but one of these did not have any associated biological processes. Where more processes were enriched, the top 6 most significant processes are shown.

| Cluster | #term ID | term description | observed<br>gene count | background<br>gene count | strength | false<br>discovery rate |
| --- | --- | --- | --- | --- | --- | --- |
| 1 | GO:0006397 | mRNA processing | 22 | 452 | 1.41 | 2.55E-22 |
|  | GO:0008380 | RNA splicing | 21 | 364 | 1.49 | 2.55E-22 |
|  | GO:0016071 | mRNA metabolic process | 23 | 613 | 1.3 | 2.01E-21 |
|  | GO:0006396 | RNA processing | 24 | 828 | 1.19 | 3.45E-20 |
|  | GO:0016070 | RNA metabolic process | 26 | 1498 | 0.97 | 4.87E-17 |
|  | GO:0000398 | mRNA splicing, via spliceosome | 15 | 231 | 1.54 | 4.87E-16 |
| 2 | GO:0007010 | Cytoskeleton organization | 33 | 1243 | 1.02 | 7.29E-23 |
|  | GO:0006996 | Organelle organization | 39 | 3482 | 0.65 | 1.48E-15 |
|  | GO:0016043 | Cellular component organization | 45 | 5482 | 0.51 | 1.01E-14 |
|  | GO:0050808 | Synapse organization | 17 | 338 | 1.3 | 3.33E-14 |
|  | GO:0007017 | Microtubule-based process | 22 | 840 | 1.02 | 9.57E-14 |
|  | GO:0000226 | Microtubule cytoskeleton organization | 19 | 559 | 1.13 | 2.01E-13 |
| 3 | GO:0031497 | Chromatin assembly | 17 | 163 | 1.78 | 6.58E-22 |
|  | GO:0051276 | Chromosome organization | 25 | 962 | 1.17 | 4.16E-21 |
|  | GO:0006325 | Chromatin organization | 21 | 599 | 1.3 | 2.2E-19 |
|  | GO:0006996 | Organelle organization | 32 | 3482 | 0.72 | 1.37E-16 |
|  | GO:0006338 | Chromatin remodeling | 15 | 284 | 1.48 | 2.82E-15 |
|  | GO:0034728 | Nucleosome organization | 12 | 143 | 1.68 | 6.33E-14 |
| 4 | GO:0110011 | Regulation of basement membrane organization | 3 | 11 | 2.82 | 0.00028 |
|  | GO:0030198 | Extracellular matrix organization | 5 | 281 | 1.64 | 0.00031 |
|  | GO:2001046 | Positive regulation of integrin-mediated signaling pathway | 3 | 15 | 2.69 | 0.00031 |
|  | GO:0007275 | Multicellular organism development | 9 | 4513 | 0.68 | 0.0016 |
|  | GO:0031589 | Cell-substrate adhesion | 4 | 197 | 1.69 | 0.0017 |
|  | GO:0009888 | Tissue development | 7 | 1902 | 0.95 | 0.0021 |
| 5 | GO:0031267 | Small GTPase binding | 4 | 288 | 1.64 | 0.0052 |
| 6 | GO:0036064 | Ciliary basal body | 3 | 181 | 2.08 | 0.0012 |
|  | GO:0000242 | Pericentriolar material | 2 | 24 | 2.78 | 0.0042 |
|  | GO:0034451 | Centriolar satellite | 2 | 109 | 2.13 | 0.0314 |

In the cytoplasmic fractions, cluster analysis identified twelve clusters in the cortex (Supplementary Figure 5), while only eight were identified in the cerebellum (Supplementary Figure 6). However, of these, five cortical and four cerebellar clusters were not associated with any gene ontology biological processes. As with the nuclear samples, there was considerable overlap in the enriched biological processes of these clusters, with cytoskeletal functions heavily featured, and synaptic pruning and the classical complement pathway enriched in both brain regions (Tables 3-4). Poly(A) RNA export from the nucleus was also enriched in both brain regions, but additional RNA export processes were only enriched in the cerebellum. Conversely, a number of synaptic functions and G protein binding functions were only enriched in the cortex, once again highlighting differences in the functional interactome of TDP-43 between the two brain regions.

**Table 3:** Top biological processes associated with each TDP-43 protein interactor in the cortex cytoplasm fraction detected using mass spectrometry analysis. Twelve clusters were identified using STRING analysis, but five of these did not have any associated biological processes. Where more processes were enriched, the top 6 most significant processes are shown.

| Cluster | #term ID | term description | observed<br>gene count | background<br>gene count | strength | false<br>discovery rate |
| --- | --- | --- | --- | --- | --- | --- |
| 1 | GO:0007010 | Cytoskeleton organization | 32 | 1243 | 0.87 | 7.24E-16 |
|  | GO:0007017 | Microtubule-based process | 27 | 840 | 0.97 | 3.21E-15 |
|  | GO:0000226 | Microtubule cytoskeleton organization | 22 | 559 | 1.05 | 1.41E-13 |
|  | GO:0006996 | Organelle organization | 44 | 3482 | 0.56 | 4.66E-13 |
|  | GO:0007399 | Nervous system development | 34 | 2190 | 0.65 | 2.59E-11 |
|  | GO:0016043 | Cellular component organization | 51 | 5482 | 0.43 | 4.93E-11 |
| 2 | GO:0007010 | Cytoskeleton organization | 8 | 1243 | 1.03 | 0.0018 |
|  | GO:0090257 | Regulation of muscle system process | 5 | 263 | 1.5 | 0.0025 |
|  | GO:0097435 | Supramolecular fiber organization | 6 | 562 | 1.25 | 0.0025 |
|  | GO:0010927 | Cellular component assembly involved in morphogenesis | 4 | 120 | 1.75 | 0.0027 |
|  | GO:0042692 | Muscle cell differentiation | 5 | 314 | 1.43 | 0.0027 |
|  | GO:0055001 | Muscle cell development | 4 | 182 | 1.57 | 0.009 |
| 3 | GO:0072583 | Clathrin-dependent endocytosis | 6 | 31 | 2.63 | 5.1E-11 |
|  | GO:0099003 | Vesicle-mediated transport in synapse | 7 | 147 | 2.02 | 7.04E-10 |
|  | GO:0006898 | Receptor-mediated endocytosis | 7 | 172 | 1.95 | 1.37E-09 |
|  | GO:0031623 | Receptor internalization | 6 | 75 | 2.24 | 1.77E-09 |
|  | GO:0098884 | Postsynaptic neurotransmitter receptor internalization | 4 | 9 | 2.99 | 0.00000005 |
|  | GO:0048488 | Synaptic vesicle endocytosis | 5 | 58 | 2.27 | 9.56E-08 |
| 4 | GO:0030007 | Cellular potassium ion homeostasis | 4 | 15 | 3.07 | 2.56E-08 |
|  | GO:0036376 | Sodium ion export across plasma membrane | 4 | 14 | 3.1 | 2.56E-08 |
|  | GO:0006883 | Cellular sodium ion homeostasis | 4 | 26 | 2.83 | 5.72E-08 |
|  | GO:1990573 | Potassium ion import across plasma membrane | 4 | 43 | 2.61 | 0.000000213 |
|  | GO:0140352 | Export from cell | 5 | 483 | 1.66 | 0.0000072 |
|  | GO:1902600 | Proton transmembrane transport | 4 | 125 | 2.15 | 0.0000072 |
| 5 | GO:0016973 | poly(A)+ mRNA export from nucleus | 3 | 23 | 2.76 | 0.00024 |
| 6 | GO:0001664 | G protein-coupled receptor binding | 3 | 333 | 1.69 | 0.0432 |
|  | GO:0031683 | G-protein beta/gamma-subunit complex binding | 2 | 25 | 2.64 | 0.0432 |
| 7 | GO:0098883 | Synapse pruning | 3 | 10 | 3.34 | 0.00000261 |
|  | GO:0006958 | Complement activation, classical pathway | 3 | 35 | 2.8 | 0.0000257 |
|  | GO:0045087 | Innate immune response | 3 | 748 | 1.47 | 0.0338 |

**Table 4:** Top biological processes associated with each TDP-43 protein interactor in the cerebellum cytoplasm fraction detected using mass spectrometry analysis. Eight clusters were identified using STRING analysis, but four of these did not have any associated biological processes. Where more processes were enriched, the top 6 most significant processes are shown.

| Cluster | #term ID | term description | observed gene count | background gene count | strength | false discovery rate |
| --- | --- | --- | --- | --- | --- | --- |
| 1 | GO:0006996 | Organelle organization | 21 | 3482 | 0.6 | 0.0000146 |
|  | GO:0016043 | Cellular component organization | 25 | 5482 | 0.48 | 0.0000146 |
|  | GO:0007010 | Cytoskeleton organization | 13 | 1243 | 0.84 | 0.0000536 |
|  | GO:0050808 | Synapse organization | 8 | 338 | 1.19 | 0.00011 |
|  | GO:0006090 | Pyruvate metabolic process | 5 | 73 | 1.66 | 0.00029 |
|  | GO:0007399 | Nervous system development | 15 | 2190 | 0.66 | 0.00043 |
| 2 | GO:0007010 | Cytoskeleton organization | 7 | 1243 | 1.14 | 0.001 |
|  | GO:0006996 | Organelle organization | 8 | 3482 | 0.75 | 0.0258 |
|  | GO:0016043 | Cellular component organization | 9 | 5482 | 0.6 | 0.0258 |
|  | GO:0051222 | Positive regulation of protein transport | 4 | 332 | 1.47 | 0.0258 |
|  | GO:0031032 | Actomyosin structure organization | 3 | 117 | 1.79 | 0.03 |
|  | GO:0010737 | Protein kinase A signaling | 2 | 15 | 2.51 | 0.0405 |
| 3 | GO:0006406 | mRNA export from nucleus | 3 | 71 | 2.12 | 0.0205 |
|  | GO:0016973 | poly(A)+ mRNA export from nucleus | 2 | 23 | 2.43 | 0.041 |
|  | GO:0046831 | Regulation of RNA export from nucleus | 2 | 14 | 2.65 | 0.041 |
| 4 | GO:0098883 | Synapse pruning | 2 | 10 | 3.34 | 0.0044 |
|  | GO:0006958 | Complement activation, classical pathway | 2 | 35 | 2.8 | 0.0148 |

### TDP-43 interacts with a number of core paraspeckle proteins

One feature of the mass spectrometry data was the identification of several core paraspeckle proteins as TDP-43 interactors. Notably, FUS and PSPC1 were identified specifically in the cortical samples. Both proteins are known to play key roles in paraspeckle function [36, 37] which has previously been associated with ALS [38]. In addition, mutations in FUS are known to cause TDP-43 negative ALS [39], making these targets of particular interest, hence they were chosen as key proteins to follow up in depth. Two other core paraspeckle proteins, NONO and SFPQ, were also identified in both the cortex and cerebellar samples, thus these proteins were also selected for follow up validation.

Consistent with the mass spectrometry analysis, co-immunoprecipitation studies in 3-month-old mouse cortex and cerebellum nuclear samples demonstrated a clear interaction of TDP-43 with all four proteins within the nuclear fraction of the cortex (Fig 3a). In addition, despite its absence in the mass spectrometry results, FUS was also pulled down in the cerebellum, along with SFPQ and NONO. Consistent with the mass spectrometry findings, PSPC1 was not present in the cerebellar pull-down, although total protein levels detected in the nuclear fraction were low (Fig 3b, d), suggesting that nuclear PSPC1 levels are markedly lower in the cerebellum compared to the cortex. No other significant differences were detected in total, nuclear or cytoplasmic expression levels of any of the paraspeckle proteins assessed (Fig 3b-e).

**Fig 3.**
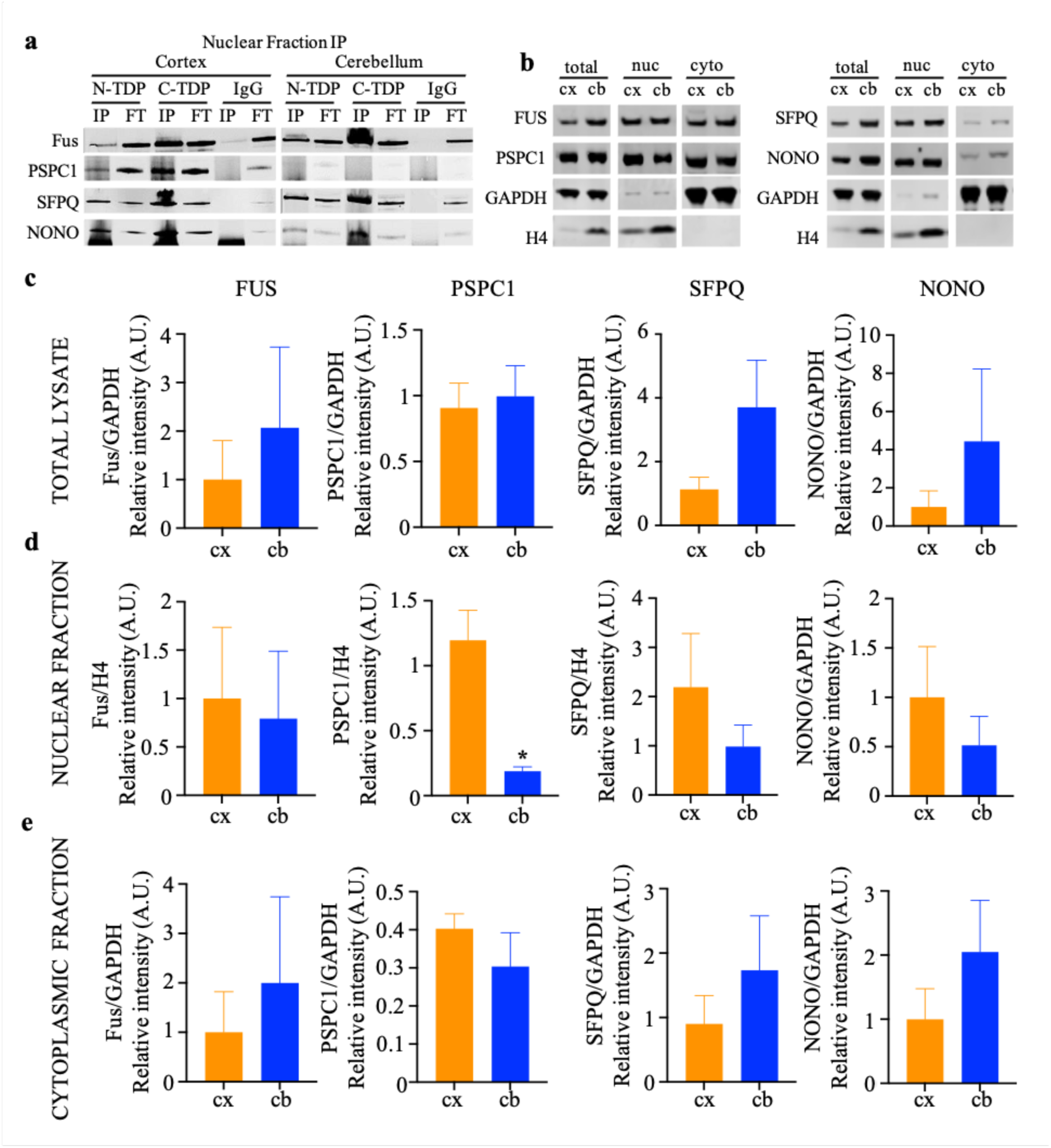
TDP-43 differentially interacts with core paraspeckle proteins FUS, PSPC1, SFPQ and NONO in the cortex and cerebellum (a) Representative examples of co-immunoprecipitation of FUS, PSPC1, SFPQ and NONO following TDP-43 pull-down in the cortex and cerebellum of 3-month-old C57Bl/6J mice with both the N (Abcam, #ab225710) and C (Proteintech #12892-1-AP) terminal TDP-43 antibodies. All four proteins were detected in the IP fraction of the cortex with both antibodies, with no band present with IgG alone. In contrast, PSPC1 was absent from the cerebellar samples following pull down with either antibody. IP: immunoprecipitation, FT: flow through. (b) Representative examples of western blot detection of total, nuclear and cytoplasmic FUS, PSPC1, SFPQ and NONO expression levels in the cortex and cerebellum of 3-month-old C57Bl/6J mice. (c-e) Quantification of FUS, PSPC1, SFPQ and NONO total (c), nuclear (d) and cytoplasmic (e) expression levels in the cortex and cerebellum of 3-month-old C57Bl/6J mice. PSPC1 levels were significantly reduced in the nuclear fraction of the cerebellum compared to the cortex. No other significant changes were observed. cx: cortex, cb: cerebellum, n=3 *p<0.05 Welch’s t-test

### TDP-43 interactions with paraspeckle proteins occur in the nucleus and cytoplasm and show region specific differences

To further validate and explore the interaction of these paraspeckle proteins with TDP-43 within the cortex and cerebellum, immunohistochemistry and proximity ligation assays were conducted with TDP-43 and each target.

As expected, immunohistochemical analysis demonstrated robust nuclear expression of all four paraspeckle proteins in the large neuronal cells of the motor cortex (Fig 4a-d). A similar staining pattern was observed in the Purkinje cells of the cerebellum for FUS, PSPC1 and SFPQ, although consistent with western blotting data, nuclear levels of PSPC1 appeared somewhat lower in the cerebellum (Fig 4b). In contrast, NONO displayed a distinct punctate pattern of staining within most large cells of the cerebellum, with puncta distributed equally throughout the nucleus and cell soma (Fig 4d). While there were clear overlaps between TDP-43 and the target paraspeckle proteins in the nucleus, there was no evidence of robust co-localisation between cytoplasmic TDP-43 and any of the proteins, despite the presence of low level cytoplasmic TDP-43 within most cells assessed.

**Fig 4.**
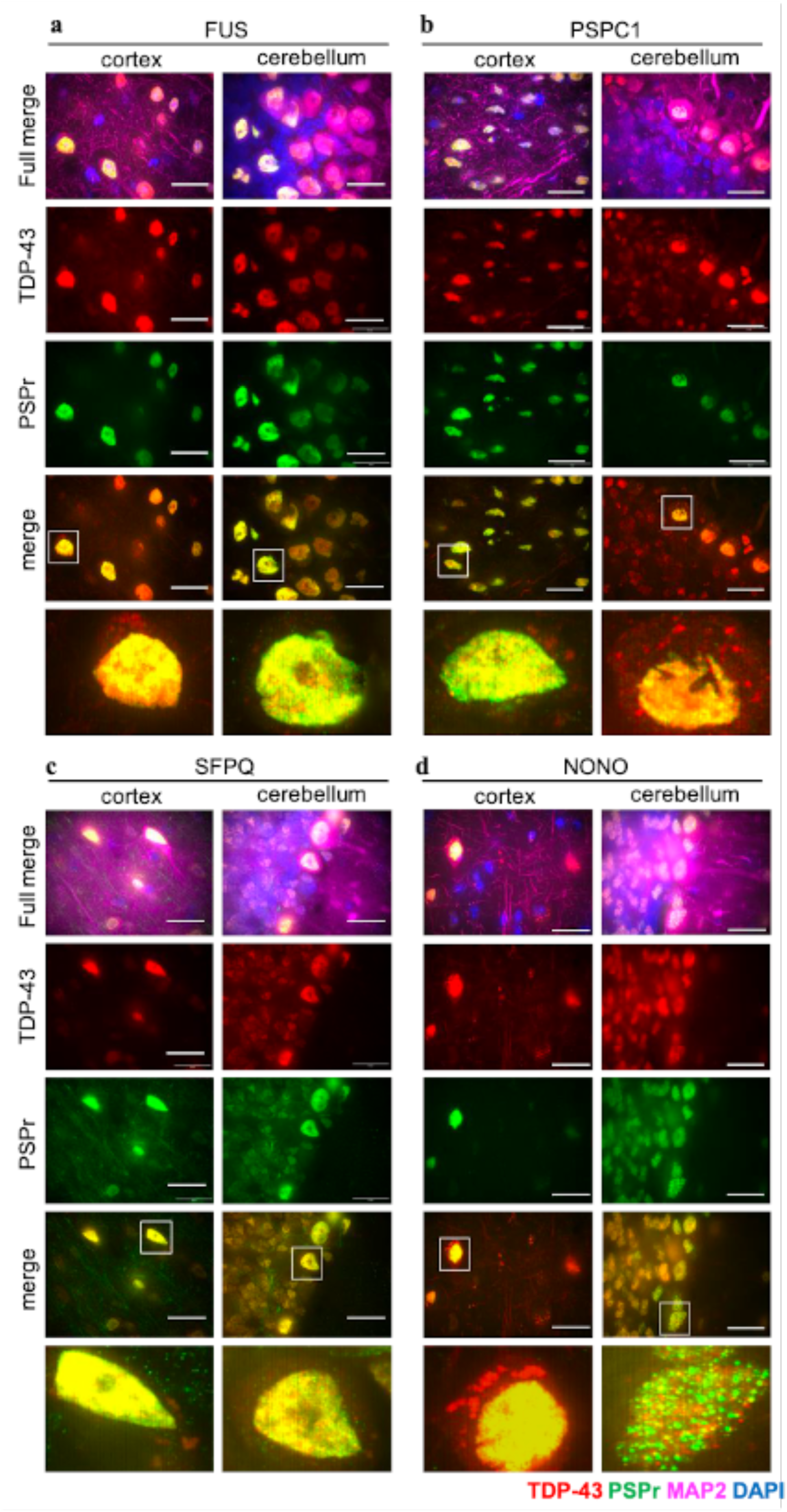
NONO shows differential cellular distribution between the cortex and cerebellum. (a-d) Representative examples of paraspeckle proteins (PSPr), FUS (a), PSPC1 (b), SFPQ (c) and NONO (d) distribution and colocalisation with TDP-43 in the motor cortex and Crus 1 region of the cerebellum. Scale bar is 20μm. All four paraspeckle proteins show abundant staining in the nucleus, with minimal evidence of expression within the cytoplasm with the exception of NONO within the cerebellum, which shows a distinct punctate pattern of staining within the nucleus and cytoplasm. TDP-43 shows abundant nuclear colocalisation with all four paraspeckle proteins in both the cortex and cerebellum, but no robust evidence of colocalization in the cytoplasm

Proximity ligation assay (PLA) analysis demonstrated interactions between TDP-43 and all four proteins in the motor cortex, and despite the absence of clear overlaps in the immunohistochemical study, these interactions were present in both the nucleus and cytoplasm of the cells (Fig 5a). The discrepancy between the PLA and IHC analysis in the cytoplasm may be due to the intense nuclear staining overwhelming the capacity to detect lower levels of the various proteins within the cytoplasm using IHC assessment. In contrast, the PLA only detects signal where an interaction occurs, hence is more likely to detect interactions where total protein levels are lower.

**Fig 5.**
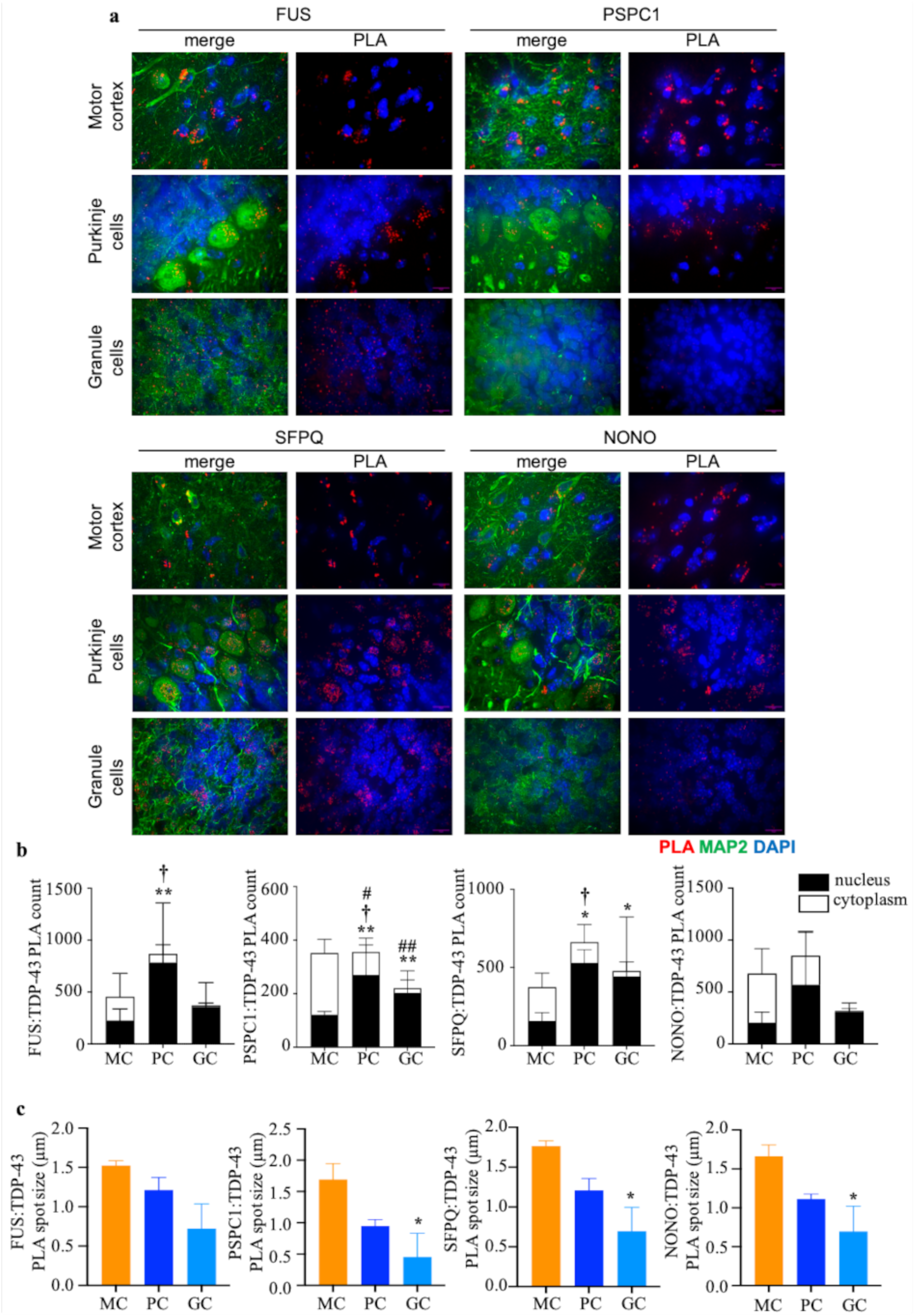
TDP-43 interactions with FUS, PSPC1, SFPQ and NONO show distinct differences between different cortical and cerebellar cell populations (a) Representative examples of proximity ligation assay (PLA) assessment of TDP-43 with FUS, PSPC1, SFPQ and NONO. (b) Quantitative analysis of PLA spot counts in the large cells of the motor cortex (MC; layer V), and the large (Purkinje cells; PC) and small (Granule cells; GC) of the cerebellum. n=3, *p<0.05; **p<0.01 cytoplasm vs. nucleus; †p<0.05 nucleus vs. motor cortex nucleus #p<0.05; ##p<0.01 cytoplasm vs. motor cortex cytoplasm. Two-way ANOVA with Fisher LSD post hoc. (c) Quantitative analysis of mean PLA spot size in large cells of the motor cortex (MC) and large (PC) and small (GC) cells of the cerebellum. n=3, *p<0.05 vs motor cortex. One way ANOVA with Tukey post hoc

PLA interactions were also detected in the cerebellum with all four interactors, including PSPC1, which had not been detected in the mass spectrometry or immunoprecipitation study. Quantification of the number of these ‘interaction’ puncta followed by two-way ANOVA analysis showed that there was a statistically significant interaction between puncta localisation and brain region for PSPC1 (p=0.0018), SFPQ (p=0.0449) and NONO (p=0.0130) with significant increases in cytoplasmic interactions of TDP-43 with PSPC1 in the cortex compared to the cerebellum (p=0.0023). FUS-TDP-43 showed a similar trend towards shifting interactions to the cytoplasm, but this did not reach significance (p=0.0954) (Supplementary Figure 9).

While these differences are linked to the different make-up of the tissue between the two brain regions, they are not associated with neuronal size *per se*. However, comparison of the large cortical neurons to both the large cells of the Purkinje layer, and the smaller cells of the granular cell layer somewhat recapitulated these findings, with a significant statistical interaction between cell type and cellular localisation detected with PSPC1 (p=0.0047). Post-hoc analysis demonstrated significantly reduced TDP-43:PSPC1 cytoplasmic interactions in the Purkinje (p=0.0274) and granule (p=0.0031) cells of the cerebellum compared to the large neurons of the motor cortex, as well as increased nuclear interactions in the Purkinje cells (p=0.0292) (Figure 5b). A similar increase in nuclear interactions in Purkinje cells compared to large motor cortex neurons was observed with SFPQ (p=0.0216). As might be expected, a significant impact of subcellular localisation was identified, with a general increase in nuclear interactions compared to cytoplasmic ones for FUS (p=0.0207), PSPC1 (p=0.0234) and SFPQ (p=0.0098). However, statistical comparison failed to detect any significant differences between the different cell populations or subcellular regions for NONO (Figure 5b).

There were also differences in the mean size of the PLA spots detected in the different cell types (Fig 5c), with spots identified in the large neurons of the motor cortex appearing larger than those of the cerebellum, reaching significance for PSPC1 (p=0.0408), SFPQ (p=0.0199) and NONO (p=0.0385) as compared to the small granule cells. FUS PLA spots showed a similar trend, although this did not reach significance (p=0.0747). These larger spots may be evidence of multiple interactions in a localised region of the cell, and in the motor cortex in particular appear to be present primarily in the perinuclear region of the cytoplasm (Figure 5a). This data hence supports the notion that increased cytoplasmic interactions of TDP-43 with these paraspeckle proteins are occurring in the soma of cortical neurons.

## Discussion

TDP-43 is a core pathogenic marker of ALS and FTD that shows a distinctly different pathological profile between disease vulnerable regions such as the frontal cortex and spinal cord and the relatively resistant cerebellum [14]. Here we demonstrate, consistent with previous studies [24], that TDP-43 load in the cerebellum in healthy control mice is higher than that seen in the cortex, most notably in the large Purkinje neurons of the cerebellum, which also show a mild increase in TDP-43 levels with age. Our data also suggests that this differential expression is driven primarily by increases in cytoplasmic levels of TDP-43. This localisation raises an interesting question regarding TDP-43 pathogenesis. If the cytoplasmic TDP-43 burden is already higher in the cerebellum, how can this brain region evade the mislocalisation and aggregation of the protein that occurs in regions such as the cortex?

While the specific cause of TDP-43 nuclear clearance and cytoplasmic aggregation in disease is still unclear, both features are found throughout vulnerable cell populations in both ALS and FTD, but are rare in cerebellar cells, even in C9orf72 cases which display abundant mutation specific pathology, in the form of p62 positive di-peptide repeat aggregates and RNA foci [15–17]. These data suggest that the function and processing of TDP-43 may differ between cells of the cerebellum and cortex and understanding more about these differences may provide insight into ALS/FTD disease mechanisms. One hypothesis is that the protein interactome of TDP-43 differs between these cell populations. To assess this, we carried out mass spectrometry analysis of the protein interacting partners of TDP-43 across the two brain regions. We identified an array of protein interactors, including many key proteins already implicated in TDP-43 function. Overall, we identified markedly more protein interactors than have previously been reported [30, 40, 41]. This may be at least in part due to the different model and methods used. Previous reports have focused on cell models and used tagged-TDP-43 to enable efficient pull-down of the protein. In this study, we focus on endogenous TDP-43 from mouse cortex and cerebellum homogenates. These samples contain the network of neurons and glia that comprise the brain coupled with the endothelial cells and pericytes that make up the vascular system, thus our dataset likely incorporates TDP-43 interactors from a spectrum of cell types. It is also important to note that we did not destroy any TDP-43:RNA interactions, hence some of our putative interactors may form RNA-dependent complexes with the protein, rather than interact directly.

Nevertheless, there are overlaps between our dataset and those previously reported, in particular with respect to enriched biological processes. STRING analysis of the TDP-43 protein interactors across our four sample sets highlights a number of expected functions, based on current understanding of TDP-43 biology. A large cluster of proteins associated with an array of RNA processing and splicing functions were identified in the nuclear fraction, consistent with previous reports. TDP-43 is known to be an RNA-binding protein, and numerous studies have demonstrated its key role in splicing. Studies have suggested that TDP-43 mediates splicing repression, protecting the transcriptome by preventing aberrant splicing, notably in ALS vulnerable cell populations such as motor neurons [42] as well as repressing cryptic exon inclusion [43–46] in a number of ALS risk genes including *UNC13A* and Stathmin 2.

Cytoskeletal associated proteins were enriched in both the nuclear and cytoplasmic fractions of both brain regions. The high abundance of many of these cytoskeletal proteins within cells means that they are often identified in proteomic studies as probable contaminants, however, TDP-43 is known to shuttle across the nuclear membrane [47] and is trafficked along the axon [48], so it is perhaps not surprising that many of these cytoskeletal components, such as the neurofilament proteins, were detected in our samples.

In general, fewer proteins were detected as TDP-43 interactors in the cytoplasmic fraction, irrespective of the brain region being assessed, and here in particular, there were clear differences between the cortex and cerebellum. A number of functions associated with synaptic function were enriched only in cortical samples, suggesting that TDP-43 may perform key functions in the synapse within this brain region. Previous studies have demonstrated that TDP-43 is present in motor neuron synapses of the mouse spinal cord [49], and ALS relevant mutations in TDP-43 lead to synaptic impairments in human induced-pluripotent stem cell derived motor neurons [50], highlighting an important functional role for TDP-43 at the synapse, that may be impacted in disease. Consistent with previous studies [49] our findings suggest the presence and function of TDP-43 at the synapse may depend on the neuron/synapse type. Understanding more about the specific role of TDP-43 at the synapse may shed light on the differential vulnerability of neurons and provide insight into potential disease mechanisms that might be targeted for therapeutic development.

Among the proteins identified as TDP-43 interactors within the cortex were a number of core paraspeckle proteins, namely SFPQ, NONO, FUS AND PSPC1. Paraspeckles are nuclear membraneless organelles comprised of NEAT1_2 long non-coding RNA and an array of RNA binding proteins; and are involved in number of cellular processes. Their formation was found to be compromised in FUS (Fused in sarcoma) linked ALS, although there was no evidence of disruption in other ALS cases [51]. TDP-43 has previously been reported to bind to and upregulate NEAT 1_2 expression [52, 53], an upregulation that is also seen in early ALS, and has been shown to have cytotoxic impacts in cell studies. Conversely, depletion of TDP-43 has also been shown to increase Neat1_2 expression and enhance paraspeckle formation in cells [54]. TDP-43 has been reported to form nuclear bodies in response to oxidative stress, that partially colocalise with *NEAT1* RNA foci. Knockdown of *NEAT1* dramatically reduces the number of these TDP-43 nuclear bodies and mutation driven inhibition of nuclear body formation during stress drives increases in cytoplasmic TDP-43 granules and is associated with increased cytotoxicity [55]. Clearly there is a complex bi-directional relationship between TDP-43 and *NEAT1* that may have profound consequences for cell viability, although some of the specifics are yet to be discerned.

However, less attention has been paid to an association between TDP-43 and the paraspeckle proteins themselves. FUS, SFPQ and NONO have been identified in the previously mentioned TDP-43 interaction studies [30, 40, 41], but PSPC1 was not specifically identified. FUS:SFPQ paraspeckle protein interactions have been reported to be disrupted in TDP-43 associated disease, although this was also seen in FUS-ALS/FTD, Progressive Supranuclear Palsy (PSP) and Corticobasal Degeneration (CBD) [56]. This is perhaps not surprising, as FUS is also a core paraspeckle protein, and TDP-43 pathology has been reported in a subset of PSP and CBD cases, which are also associated with SFPQ depletion [57].

Intriguingly, our data suggests that TDP-43:paraspeckle protein interactions may not be confined to the nucleus, with cytoplasmic interactions detected in all cell populations assessed, but enriched in large cortical neurons within the motor cortex compared to Purkinje and granule cells within the cerebellum. Studies in HEK293 cell models have shown that SFPQ and NONO co-aggregate with TDP-43 in the cytoplasm in a model of TDP-43 aggregation [58], while osmotic stress can shift the localisation of SFPQ, NONO and PSPC1 to the cytoplasm, where they colocalise and form large granules [59]. This cytoplasmic accumulation is not typically associated with Neat1_2, which forms distinct cytoplasmic puncta. *NEAT1* expression has also been observed at low levels in the mono-ribosomal and cytoplasmic fractions of non-stressed cells [60], and cytoplasmic Neat1_1 has been found to modify glycolysis in breast cancer, via an interaction with the PGK1, PGAM1 and ENO1 enzymes [61]. While this interaction involves Neat1_1, rather than the longer Neat1_2 isoform that has been linked to paraspeckle formation, it nevertheless demonstrates the potential for robust cytoplasmic interactions and roles of these RNAs.

Our current data suggests that cytoplasmic TDP-43:paraspeckle protein interactions occur within the cytoplasm of healthy neurons, and these interactions are enhanced in the disease vulnerable cortex, compared to the relatively resistant cerebellum. This finding is particularly robust for PSPC1, which has been shown to display persistent fibril formation in response to osmotic stress, that differentiates it from SFPQ and NONO [59]. This raises the question of whether alterations in this TDP-43 interaction could play a key role in the onset and progression of TDP-43 linked pathogenesis. It is important to note that our current data does not elucidate whether these cytoplasmic interactions are direct, or via interactions with *NEAT1*, clearly a key question that needs to be addressed. Similarly, the functional role of these TDP-43:paraspeckle protein interactions in the cytoplasm requires elucidation. Further investigation into what these complexes do, and whether they are altered in TDP-43 pathogenesis could provide insight into disease processes in sporadic disease and highlight potential novel targets for targeted intervention to prevent further disease progression.

In summary, we have confirmed that TDP-43 protein levels are increased within the cerebellum compared to the cortex and demonstrated that this increased protein burden appears to be primarily driven by an increase in cytoplasmic load. We have identified a number of core differences in the TDP-43 interactome between these two brain regions, notably the core paraspeckle protein, PSPC1, and have demonstrated that the localisation and frequency of TDP-43:paraspeckle protein interactions varies according to cell type/brain region, with increased cytoplasmic interactions identified in cortical neurons compared to those in the cerebellum. Further investigation into these cellular/regional differences in TDP-43 interactors may improve our understanding of disease processes and lead to new therapeutic approaches.

## Declarations

### Consent for publication

Not applicable

### Availability of data and materials

The mass spectrometry proteomics dataset generated during the current study has been deposited to the ProteomeXchange Consortium via the PRIDE partner repository (http://www.ebi.ac.uk/pride) with the dataset identifier PXD062532 and 10.6019/PXD062532. All other data generated or analysed during this study are included in this published article and its supplementary information files.

### Author Contributions

TB and ST conducted the experimental work, with the guidance of CV and JM. SL carried out the mass spectrometry work and basic analysis. TB and JM carried out the remainder of the analysis and constructed the figures. JM wrote the manuscript. All authors reviewed and edited the manuscript.

## Acknowledgements

This work was supported by The John and Lucille Van Geest Foundation and the Motor Neurone Disease Association

## References

1. Neumann, M., et al., Ubiquitinated TDP-43 in frontotemporal lobar degeneration and amyotrophic lateral sclerosis. Science, 2006. 314(5796): p. 130–3.

2. Arai, T., et al., TDP-43 is a component of ubiquitin-positive tau-negative inclusions in frontotemporal lobar degeneration and amyotrophic lateral sclerosis. Biochem Biophys Res Commun, 2006. 351(3): p. 602–11.

3. Sreedharan, J., et al., TDP-43 mutations in familial and sporadic amyotrophic lateral sclerosis. Science, 2008. 319(5870): p. 1668–72.

4. Gitcho, M.A., et al., TDP-43 A315T mutation in familial motor neuron disease. Ann Neurol, 2008. 63(4): p. 535–8.

5. Kabashi, E., et al., TARDBP mutations in individuals with sporadic and familial amyotrophic lateral sclerosis. Nat Genet, 2008. 40(5): p. 572–4.

6. Murray, M.E., et al., Clinical and neuropathologic heterogeneity of c9FTD/ALS associated with hexanucleotide repeat expansion in C9ORF72. Acta Neuropathol, 2011. 122(6): p. 673–90.

7. Smith, B.N., et al., Mutations in the vesicular trafficking protein annexin A11 are associated with amyotrophic lateral sclerosis. Sci Transl Med, 2017. 9(388).

8. Kamada, M., et al., Clinicopathologic features of autosomal recessive amyotrophic lateral sclerosis associated with optineurin mutation. Neuropathology, 2014. 34(1): p. 64–70.

9. Geser, F., et al., Evidence of multisystem disorder in whole-brain map of pathological TDP-43 in amyotrophic lateral sclerosis. Arch Neurol, 2008. 65(5): p. 636–41.

10. Kawles, A., et al., Cortical and subcortical pathological burden and neuronal loss in an autopsy series of FTLD-TDP-type C. Brain, 2022. 145(3): p. 1069–1078.

11. Daskalakis, Z.J., et al., Exploring the connectivity between the cerebellum and motor cortex in humans. J Physiol, 2004. 557(Pt 2): p. 689–700.

12. Allen, G., et al., Magnetic resonance imaging of cerebellar-prefrontal and cerebellar-parietal functional connectivity. Neuroimage, 2005. 28(1): p. 39–48.

13. Van Overwalle, F. and P. Marien, Functional connectivity between the cerebrum and cerebellum in social cognition: A multi-study analysis. Neuroimage, 2016. 124(Pt A): p. 248–255.

14. Kawakami, I., T. Arai, and M. Hasegawa, The basis of clinicopathological heterogeneity in TDP-43 proteinopathy. Acta Neuropathol, 2019. 138(5): p. 751–770.

15. Troakes, C., et al., An MND/ALS phenotype associated with C9orf72 repeat expansion: abundant p62-positive, TDP-43-negative inclusions in cerebral cortex, hippocampus and cerebellum but without associated cognitive decline. Neuropathology, 2012. 32(5): p. 505–14.

16. Mehta, A.R., et al., Improved detection of RNA foci in C9orf72 amyotrophic lateral sclerosis post-mortem tissue using BaseScope shows a lack of association with cognitive dysfunction. Brain Commun, 2020. 2(1): p. fcaa009.

17. Al-Sarraj, S., et al., p62 positive, TDP-43 negative, neuronal cytoplasmic and intranuclear inclusions in the cerebellum and hippocampus define the pathology of C9orf72-linked FTLD and MND/ALS. Acta Neuropathol, 2011. 122(6): p. 691–702.

18. Kaliszewska, A., et al., Elucidating the Role of Cerebellar Synaptic Dysfunction in C9orf72-ALS/FTD - a Systematic Review and Meta-Analysis. Cerebellum, 2022. 21(4): p. 681–714.

19. Bede, P., et al., Genotype-associated cerebellar profiles in ALS: focal cerebellar pathology and cerebro-cerebellar connectivity alterations. J Neurol Neurosurg Psychiatry, 2021. 92(11): p. 1197–1205.

20. Pizzarotti, B., et al., Frontal and Cerebellar Atrophy Supports FTSD-ALS Clinical Continuum. Front Aging Neurosci, 2020. 12: p. 593526.

21. Tan, R.H., et al., Cerebellar neuronal loss in amyotrophic lateral sclerosis cases with ATXN2 intermediate repeat expansions. Ann Neurol, 2016. 79(2): p. 295–305.

22. Brettschneider, J., et al., TDP-43 pathology and neuronal loss in amyotrophic lateral sclerosis spinal cord. Acta Neuropathol, 2014. 128(3): p. 423–37.

23. Yousef, A., et al., Neuron loss and degeneration in the progression of TDP-43 in frontotemporal lobar degeneration. Acta Neuropathol Commun, 2017. 5(1): p. 68.

24. Hu, W., et al., Expression of Tau Pathology-Related Proteins in Different Brain Regions: A Molecular Basis of Tau Pathogenesis. Front Aging Neurosci, 2017. 9: p. 311.

25. Polymenidou, M., et al., Misregulated RNA processing in amyotrophic lateral sclerosis. Brain Res, 2012. 1462: p. 3–15.

26. Tollervey, J.R., et al., Characterizing the RNA targets and position-dependent splicing regulation by TDP-43. Nat Neurosci, 2011. 14(4): p. 452–8.

27. Xiao, S., et al., RNA targets of TDP-43 identified by UV-CLIP are deregulated in ALS. Mol Cell Neurosci, 2011. 47(3): p. 167–80.

28. Aulas, A. and C. Vande Velde, Alterations in stress granule dynamics driven by TDP-43 and FUS: a link to pathological inclusions in ALS? Front Cell Neurosci, 2015. 9: p. 423.

29. Diaper, D.C., et al., Drosophila TDP-43 dysfunction in glia and muscle cells cause cytological and behavioural phenotypes that characterize ALS and FTLD. Hum Mol Genet, 2013. 22(19): p. 3883–93.

30. Freibaum, B.D., et al., Global analysis of TDP-43 interacting proteins reveals strong association with RNA splicing and translation machinery. J Proteome Res, 2010. 9(2): p. 1104–20.

31. Davis, S.A., et al., TDP-43 interacts with mitochondrial proteins critical for mitophagy and mitochondrial dynamics. Neurosci Lett, 2018. 678: p. 8–15.

32. Volkening, K., et al., RNA and Protein Interactors with TDP-43 in Human Spinal-Cord Lysates in Amyotrophic Lateral Sclerosis. J Proteome Res, 2018. 17(4): p. 1712–1729.

33. Eng, J.K., A.L. McCormack, and J.R. Yates, An approach to correlate tandem mass spectral data of peptides with amino acid sequences in a protein database. J Am Soc Mass Spectrom, 1994. 5(11): p. 976–89.

34. Perez-Riverol, Y., et al., The PRIDE database at 20 years: 2025 update. Nucleic Acids Res, 2025. 53(D1): p. D543–D553.

35. Szklarczyk, D., et al., STRING v10: protein-protein interaction networks, integrated over the tree of life. Nucleic Acids Res, 2015. 43(Database issue): p. D447–52.

36. Naganuma, T., et al., Alternative 3’-end processing of long noncoding RNA initiates construction of nuclear paraspeckles. EMBO J, 2012. 31(20): p. 4020–34.

37. Fox, A.H., C.S. Bond, and A.I. Lamond, P54nrb forms a heterodimer with PSP1 that localizes to paraspeckles in an RNA-dependent manner. Mol Biol Cell, 2005. 16(11): p. 5304–15.

38. An, H., N.G. Williams, and T.A. Shelkovnikova, NEAT1 and paraspeckles in neurodegenerative diseases: A missing lnc found? Noncoding RNA Res, 2018. 3(4): p. 243–252.

39. Vance, C., et al., Mutations in FUS, an RNA processing protein, cause familial amyotrophic lateral sclerosis type 6. Science, 2009. 323(5918): p. 1208–1211.

40. Ling, S.C., et al., ALS-associated mutations in TDP-43 increase its stability and promote TDP-43 complexes with FUS/TLS. Proc Natl Acad Sci U S A, 2010. 107(30): p. 13318–23.

41. Blokhuis, A.M., et al., Comparative interactomics analysis of different ALS-associated proteins identifies converging molecular pathways. Acta Neuropathol, 2016. 132(2): p. 175–196.

42. Donde, A., et al., Splicing repression is a major function of TDP-43 in motor neurons. Acta Neuropathol, 2019. 138(5): p. 813–826.

43. Ling, J.P., et al., TDP-43 repression of nonconserved cryptic exons is compromised in ALS-FTD. Science, 2015. 349(6248): p. 650–5.

44. Melamed, Z., et al., Premature polyadenylation-mediated loss of stathmin-2 is a hallmark of TDP-43-dependent neurodegeneration. Nat Neurosci, 2019. 22(2): p. 180–190.

45. Klim, J.R., et al., ALS-implicated protein TDP-43 sustains levels of STMN2, a mediator of motor neuron growth and repair. Nat Neurosci, 2019. 22(2): p. 167–179.

46. Ma, X.R., et al., TDP-43 represses cryptic exon inclusion in the FTD-ALS gene UNC13A. Nature, 2022. 603(7899): p. 124–130.

47. Ayala, Y.M., et al., Structural determinants of the cellular localization and shuttling of TDP-43. J Cell Sci, 2008. 121(Pt 22): p. 3778–85.

48. Fallini, C., G.J. Bassell, and W. Rossoll, The ALS disease protein TDP-43 is actively transported in motor neuron axons and regulates axon outgrowth. Hum Mol Genet, 2012. 21(16): p. 3703–18.

49. Broadhead, M.J., et al., Synaptic expression of TAR-DNA-binding protein 43 in the mouse spinal cord determined using super-resolution microscopy. Front Mol Neurosci, 2023. 16: p. 1027898.

50. Lepine, S., et al., Homozygous ALS-linked mutations in TARDBP/TDP-43 lead to hypoactivity and synaptic abnormalities in human iPSC-derived motor neurons. iScience, 2024. 27(3): p. 109166.

51. Shelkovnikova, T.A., et al., Compromised paraspeckle formation as a pathogenic factor in FUSopathies. Hum Mol Genet, 2014. 23(9): p. 2298–312.

52. Nishimoto, Y., et al., The long non-coding RNA nuclear-enriched abundant transcript 1_2 induces paraspeckle formation in the motor neuron during the early phase of amyotrophic lateral sclerosis. Mol Brain, 2013. 6: p. 31.

53. Suzuki, H., et al., C9-ALS/FTD-linked proline-arginine dipeptide repeat protein associates with paraspeckle components and increases paraspeckle formation. Cell Death Dis, 2019. 10(10): p. 746.

54. Shelkovnikova, T.A., et al., Protective paraspeckle hyper-assembly downstream of TDP-43 loss of function in amyotrophic lateral sclerosis. Mol Neurodegener, 2018. 13(1): p. 30.

55. Wang, C., et al., Stress Induces Dynamic, Cytotoxicity-Antagonizing TDP-43 Nuclear Bodies via Paraspeckle LncRNA NEAT1-Mediated Liquid-Liquid Phase Separation. Mol Cell, 2020. 79(3): p. 443–458 e7.

56. Ishigaki, S., et al., Aberrant interaction between FUS and SFPQ in neurons in a wide range of FTLD spectrum diseases. Brain, 2020. 143(8): p. 2398–2405.

57. Riku, Y., et al., Motor neuron TDP-43 proteinopathy in progressive supranuclear palsy and corticobasal degeneration. Brain, 2022. 145(8): p. 2769–2784.

58. Dammer, E.B., et al., Coaggregation of RNA-binding proteins in a model of TDP-43 proteinopathy with selective RGG motif methylation and a role for RRM1 ubiquitination. PLoS One, 2012. 7(6): p. e38658.

59. Yucel-Polat, A., et al., Dynamic Localization of Paraspeckle Components under Osmotic Stress. Noncoding RNA, 2024. 10(2).

60. van Heesch, S., et al., Extensive localization of long noncoding RNAs to the cytosol and mono- and polyribosomal complexes. Genome Biol, 2014. 15(1): p. R6.

61. Park, M.K., et al., NEAT1 is essential for metabolic changes that promote breast cancer growth and metastasis. Cell Metab, 2021. 33(12): p. 2380–2397 e9.

