## Supplementary data for "Differential TDP-43 interactomes between the cortex and cerebellum in the mouse"

Supplementary Figure 1

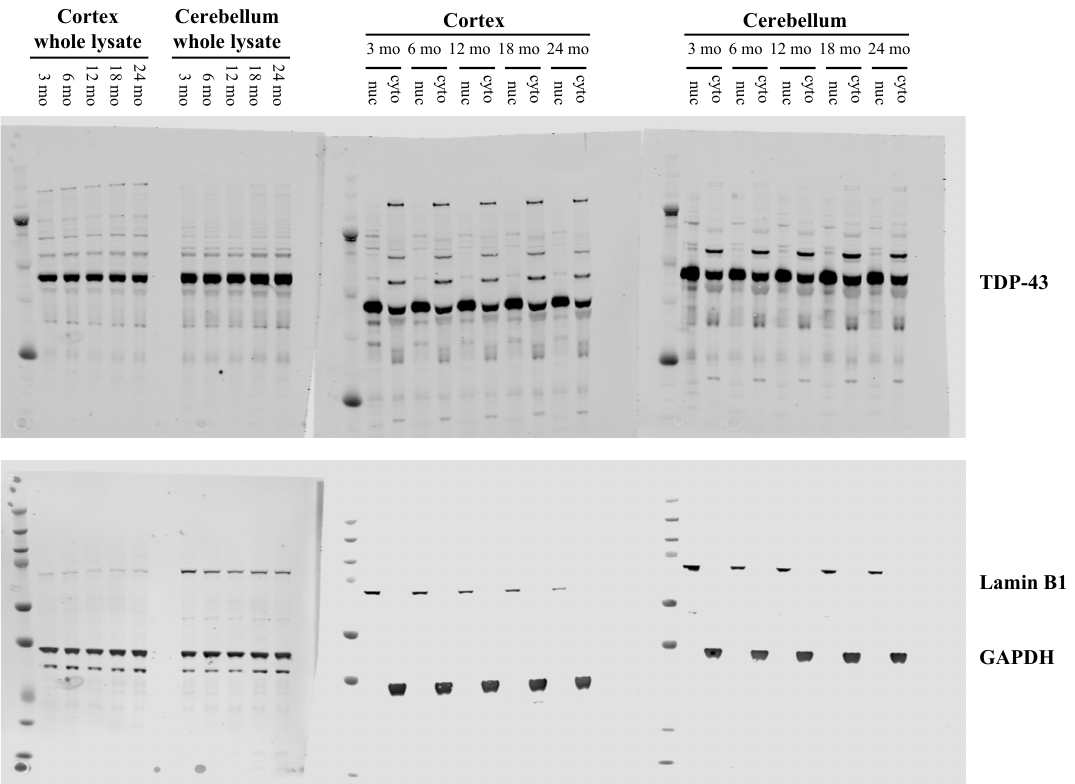

Supplementary Figure 1: Full western blots of data shown in figure 1a. Blots were probed with rabbit anti-TDP (upper) and mouse anti Lamin B1 and GAPDH (lower), and then with goat anti-rabbit Dylight 680 and goat anti-mouse Dylight 800.

Supplementary Figure 2

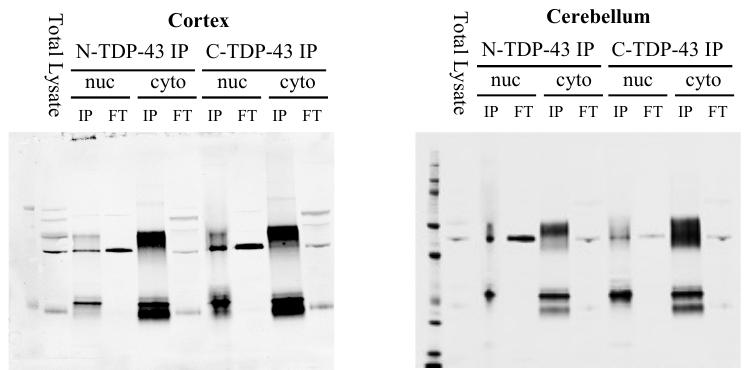

Supplementary Figure 2: Full western blots of data shown in figure 2a.

Supplementary table 1: TDP-43 epitopes identified using mass spectrometry in each sample tested.

Mass spectrometry detection of TDP-43 assigned epitopes was assessed in all samples, each consisting of one male and one female cortex or cerebellum. Identification of a minimum of two epitopes, typically > 4, was detected following C-terminal TDP-43 pull down (Proteintech #12892-1-AP). Epitope identification with the N terminal TDP-43 pull down (Abcam, #ab225710) was less robust, with some samples failing to detect a single epitope.

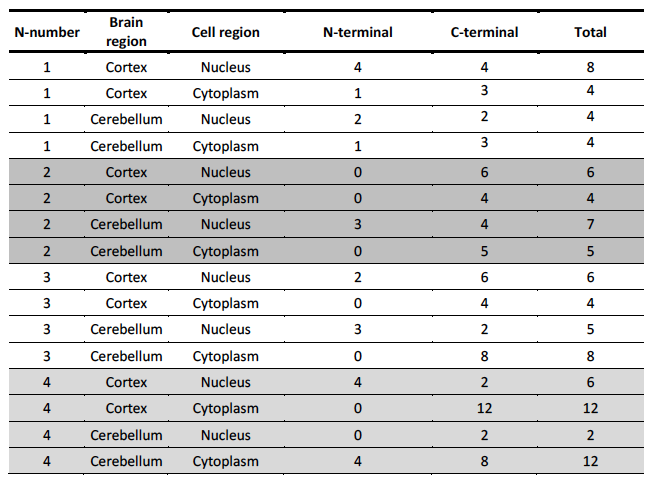

Supplementary table 2: Filtered list of TDP-43 interactors (at least two peptides in 2 different animals). Proteins are listed in alphabetical order, and their presence in a specific brain region/fraction is indicated by a ‘y’

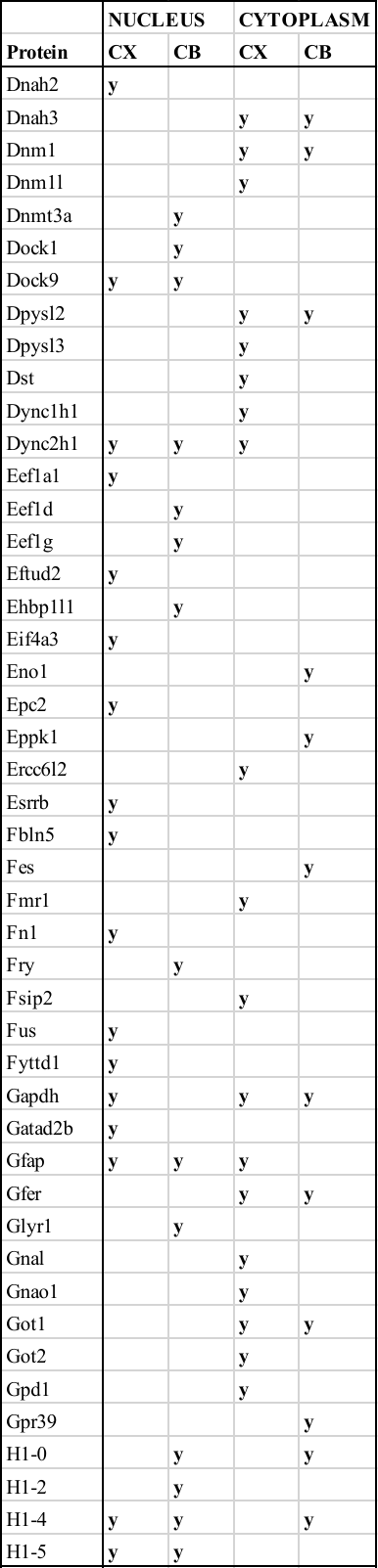

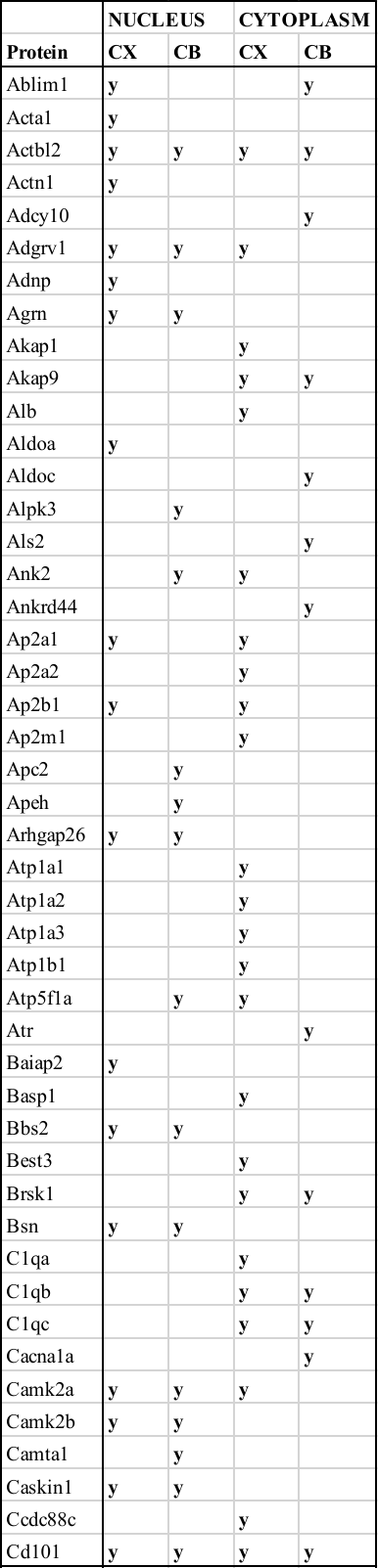

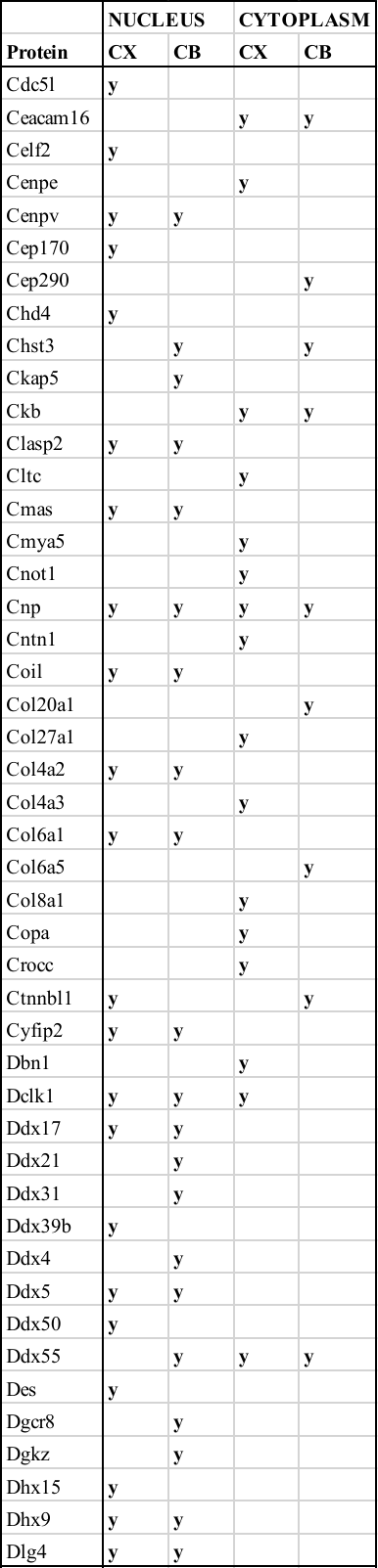

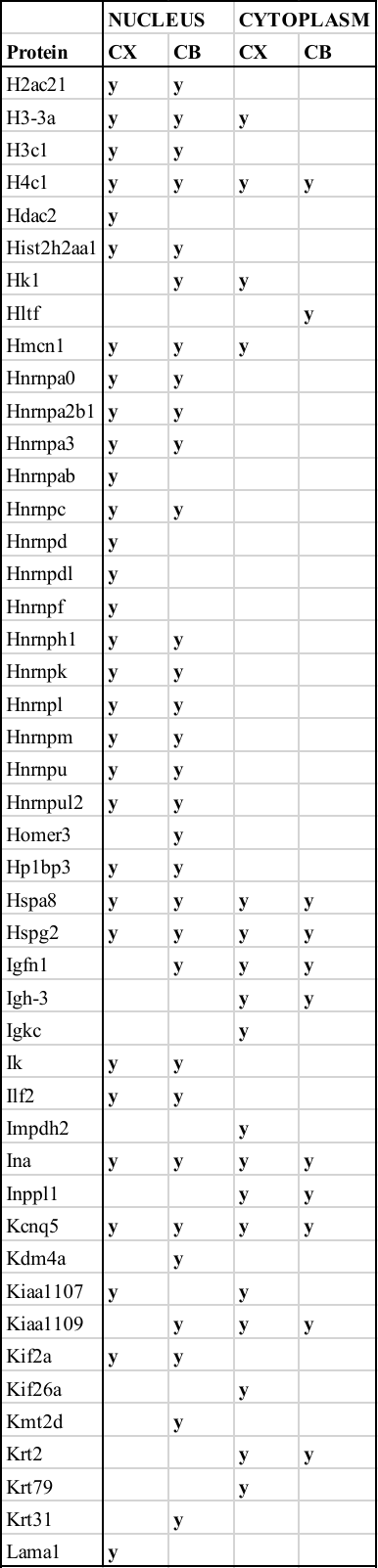

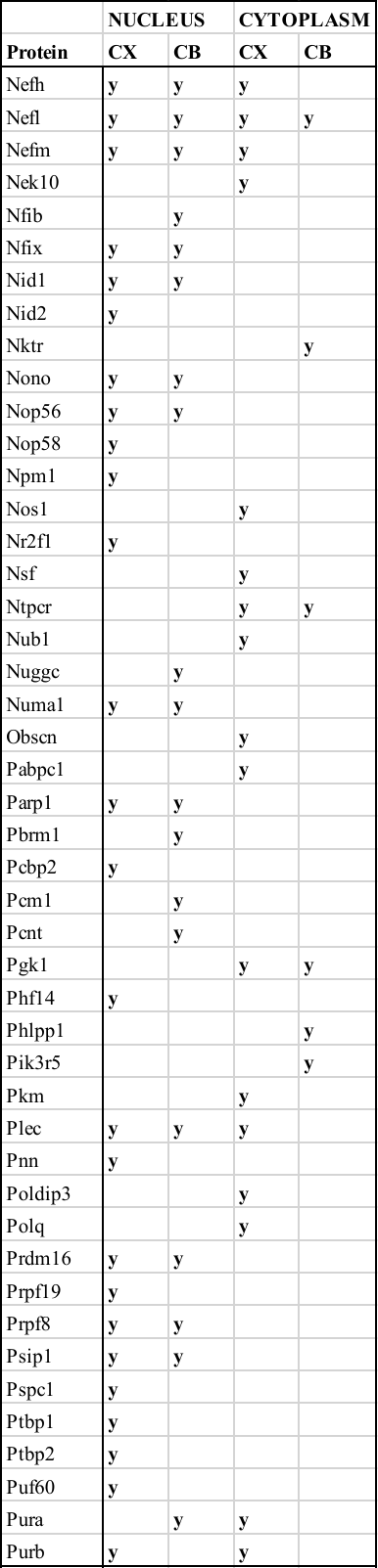

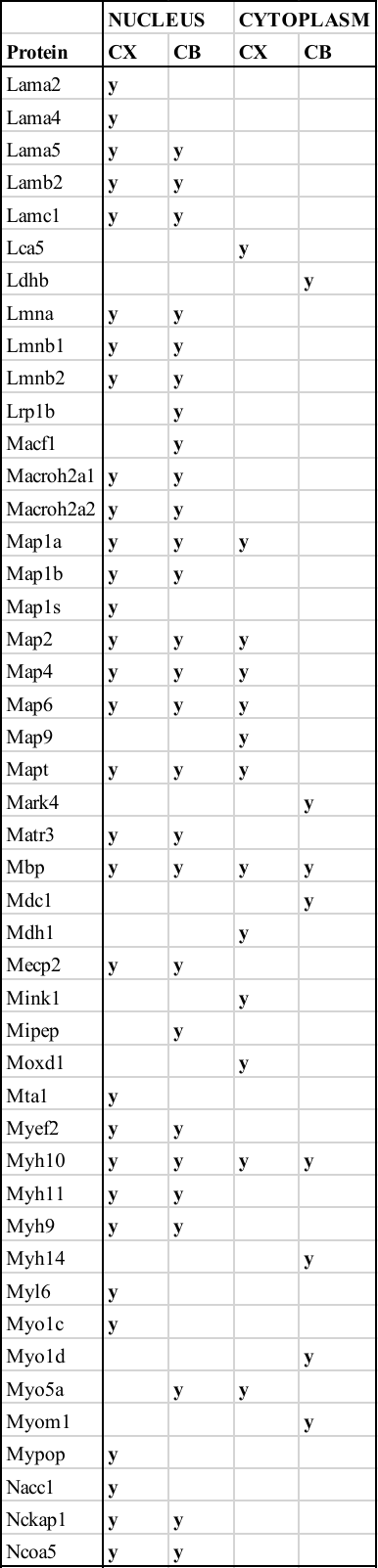

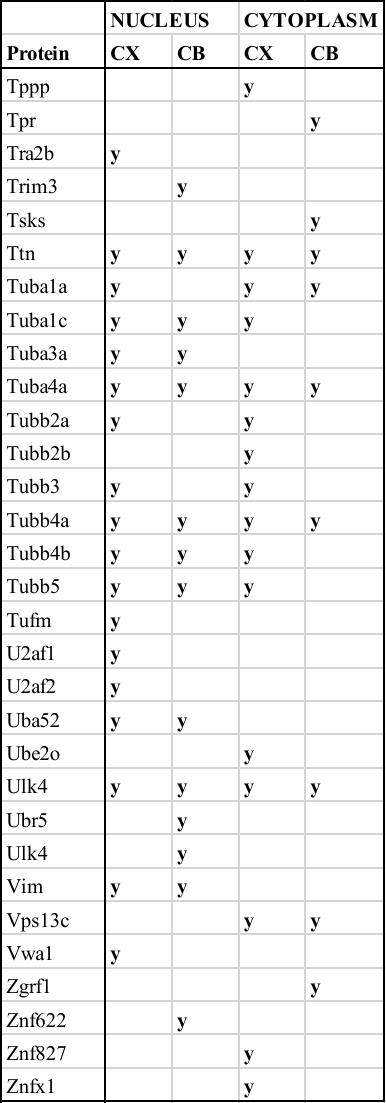

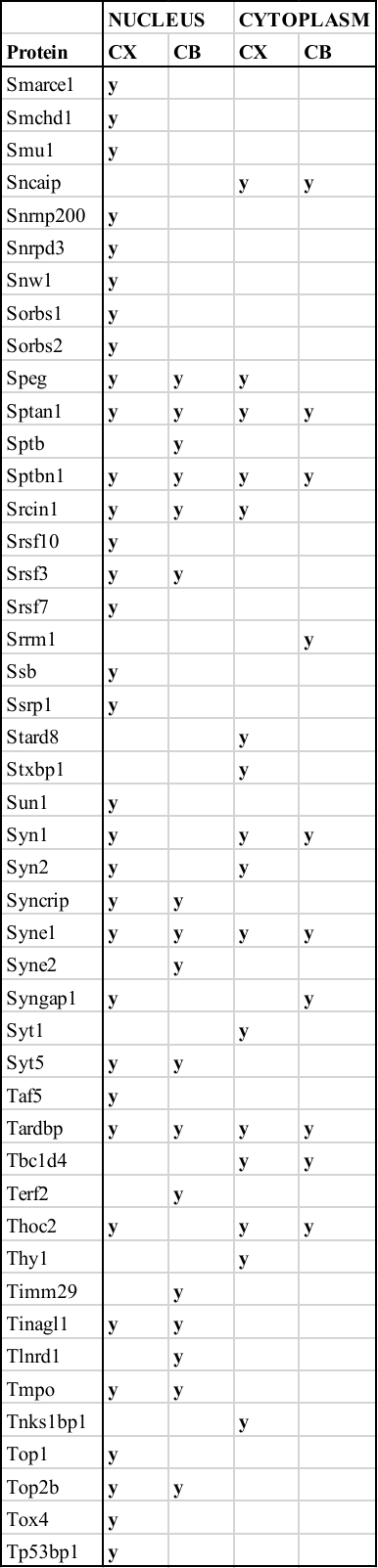

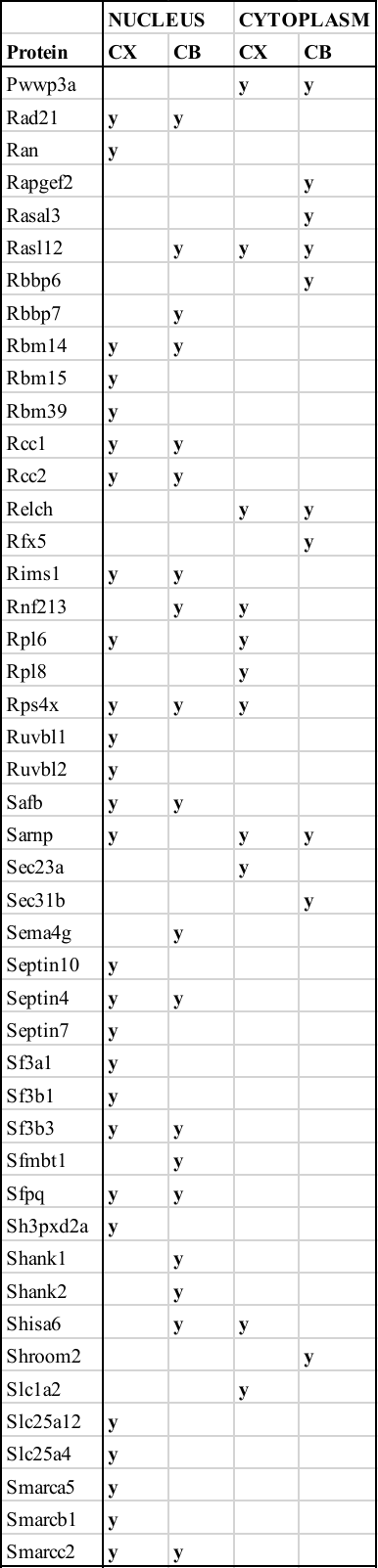

Supplementary Figure 3: STRING analysis and functional (MCL) clustering of TDP-43 protein interactors in the nuclear fraction of the cortex. Six functional clusters were identified.

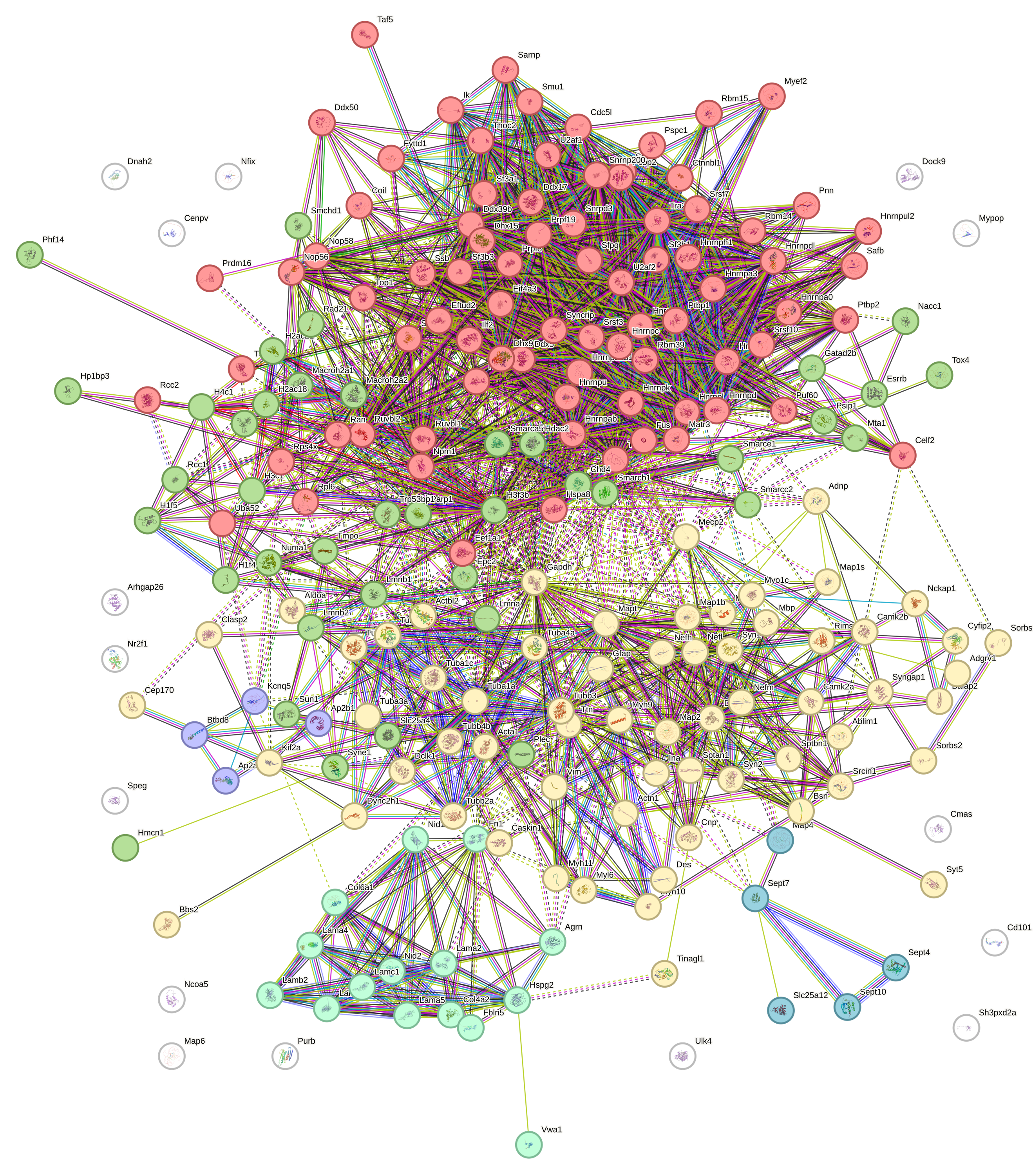

Supplementary Figure 4: STRING analysis and functional (MCL) clustering of TDP-43 protein interactors in the nuclear fraction of the cerebellum. Seven functional clusters were identified

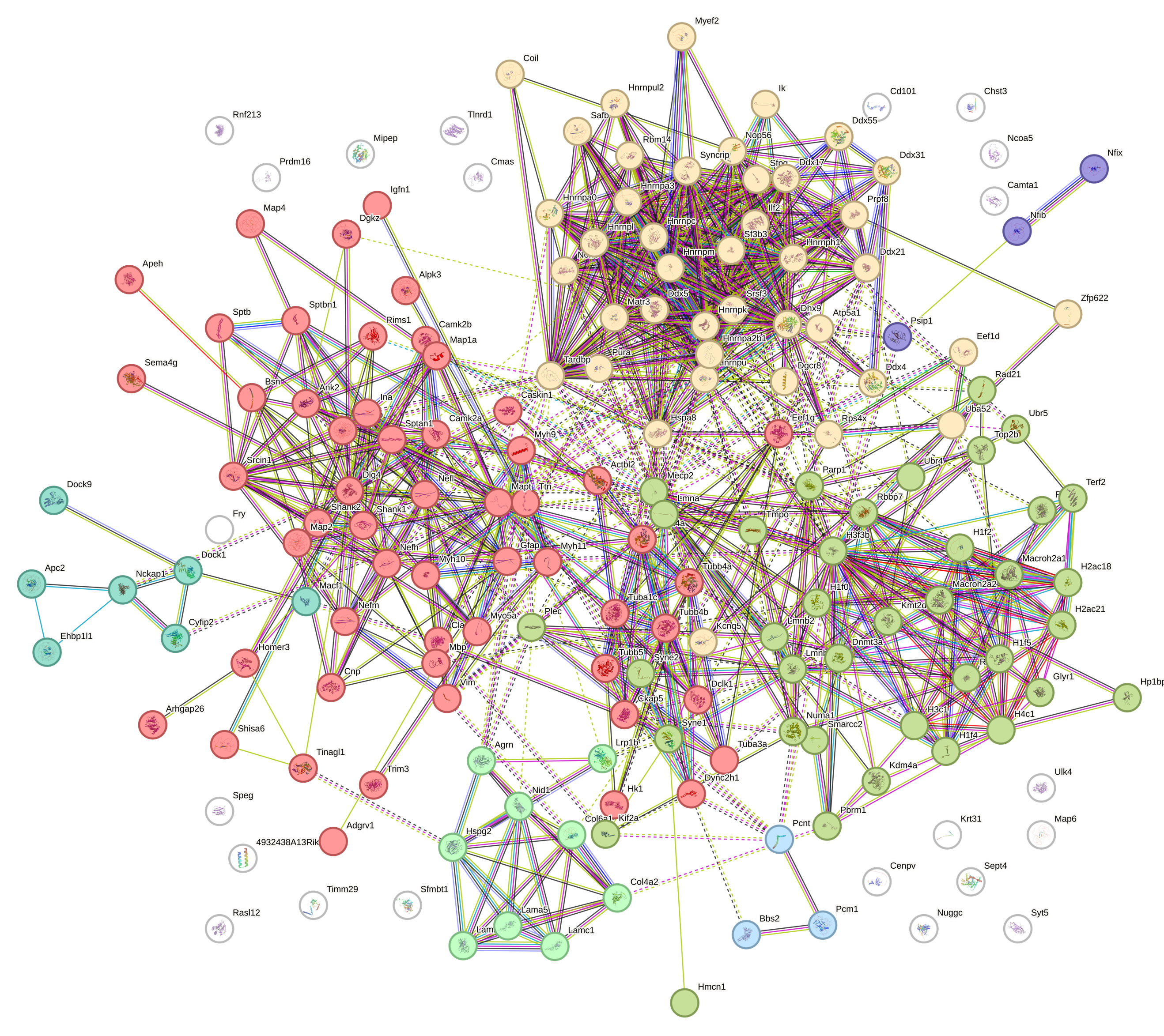

Supplementary Figure 5: STRING analysis and functional (MCL) clustering of TDP-43 protein interactors in the cytoplasmic fraction of the cortex. Twelve functional clusters were identified.

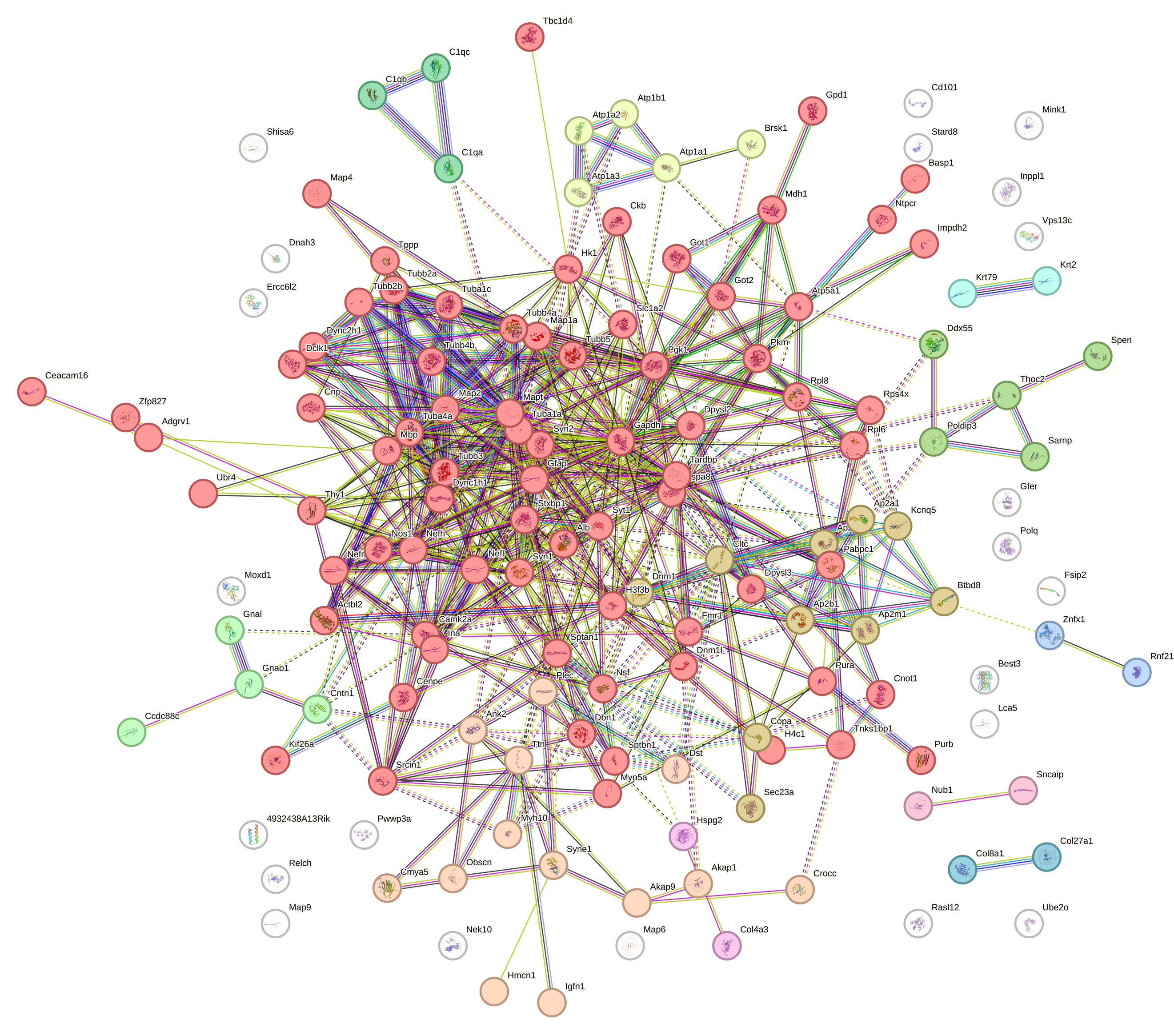

Supplementary Figure 6: STRING analysis and functional (MCL) clustering of TDP-43 protein interactors in the cytoplasmic fraction of the cerebellum. Eight functional clusters were identified.

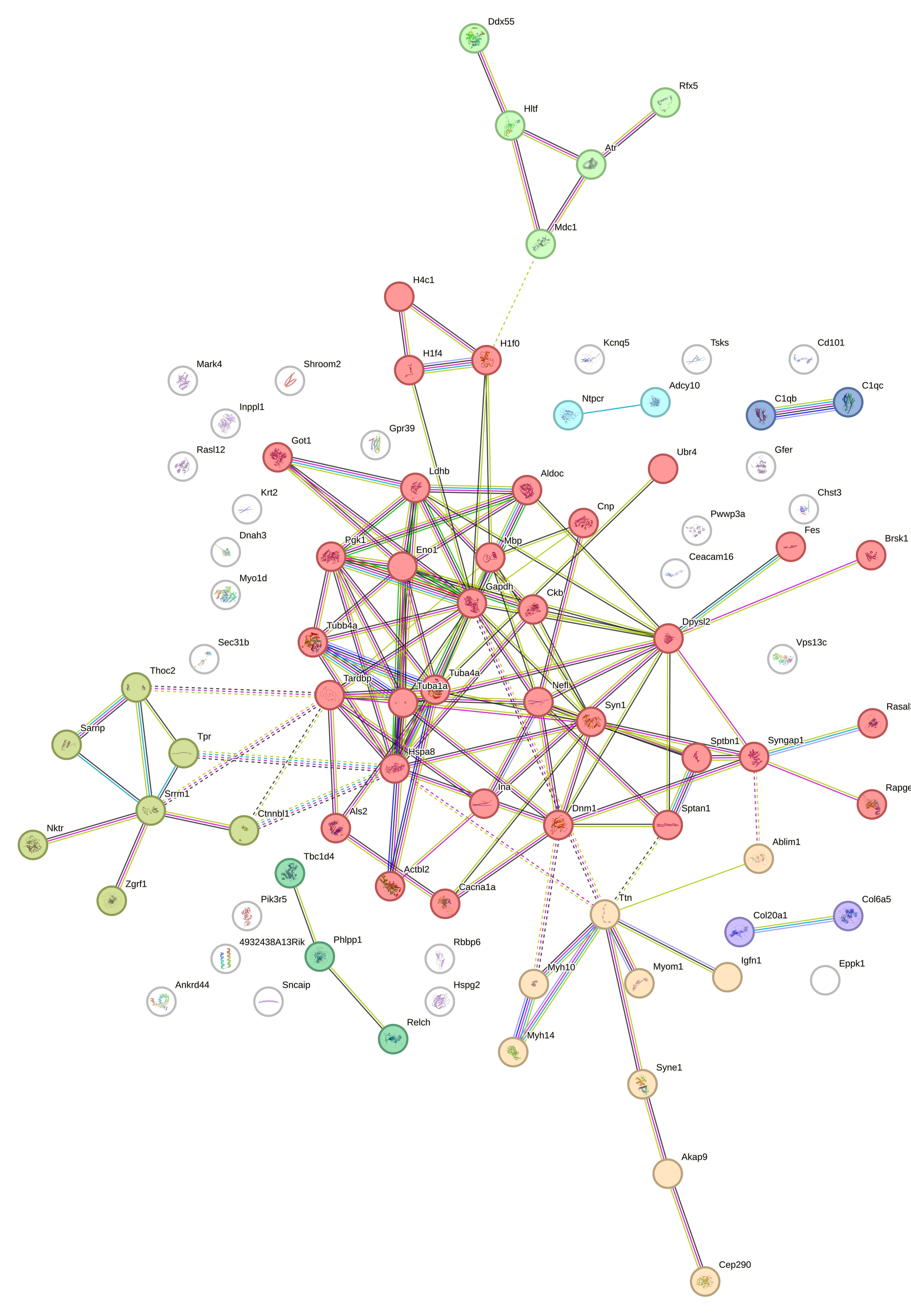

Supplementary Figure 7

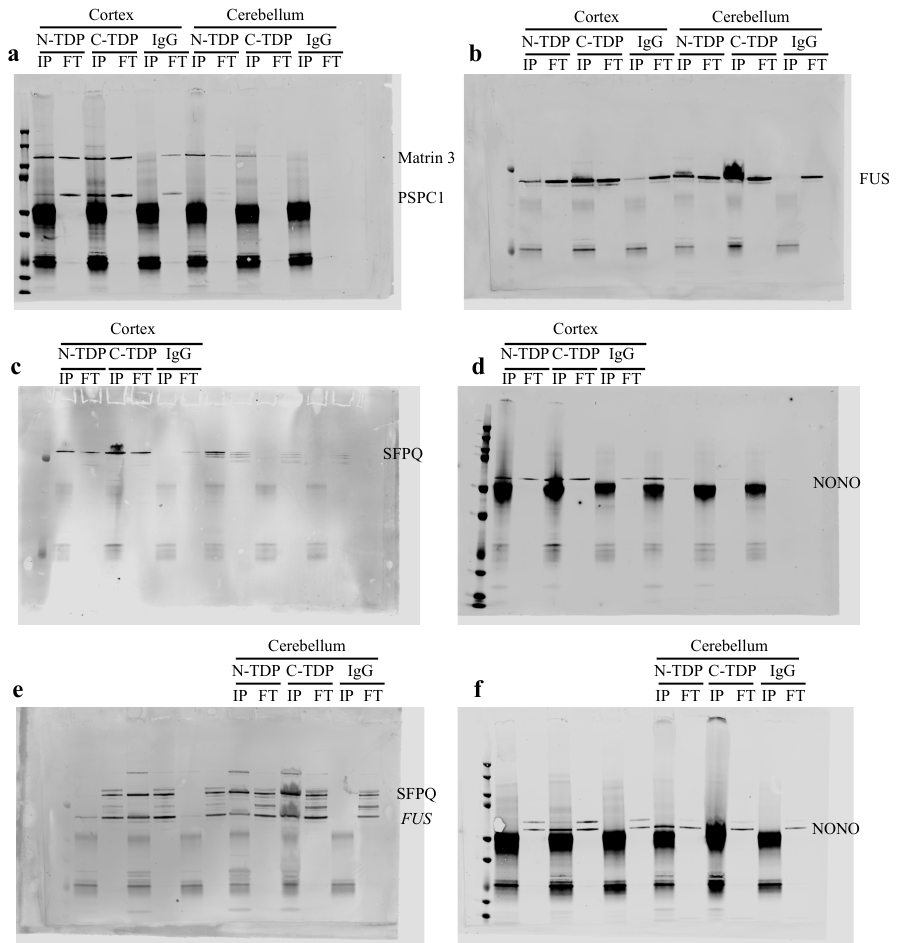

Supplementary Figure 7: Full western blot images of data shown in figure 3a. Cortex and cerebellum samples are on the same blots for PSPC1 (a) and FUS (b). Cortex and Cerebellum examples were from different gels for SFPQ (c, e) and NONO (d, f). Note Matrin 3 (not reported in this study) was also probed in (a), and FUS was additionally probed in (e).

Supplementary Figure 8

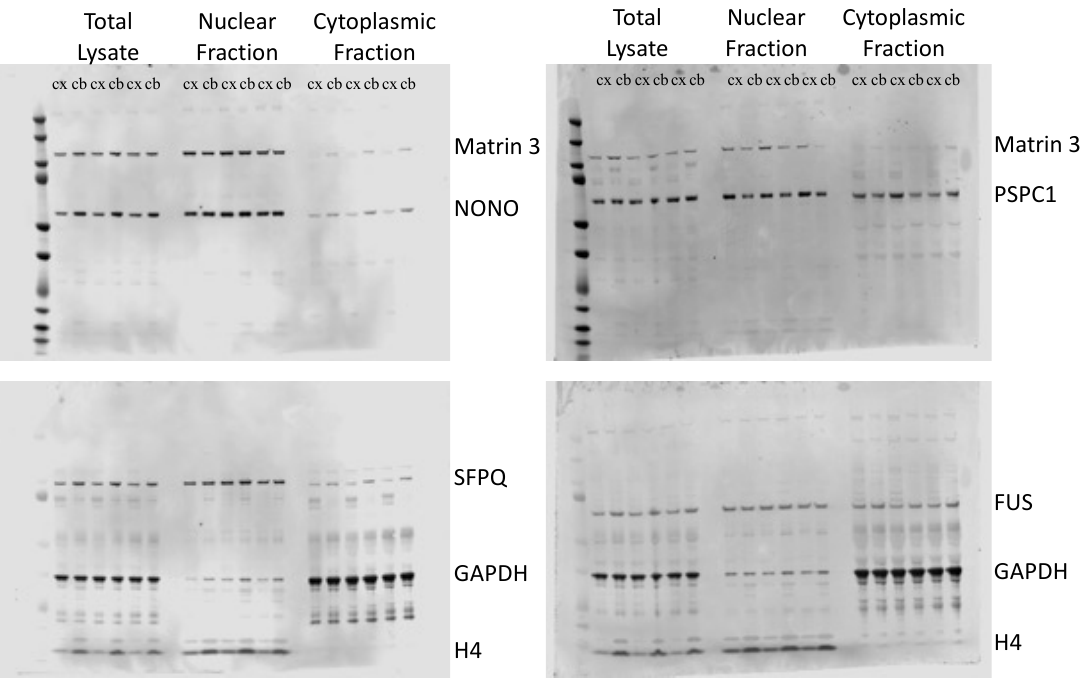

Supplementary Figure 8: Full western blot images of data shown in figure 3b. Blots were probed with several primary antibodies as noted on the blot (note Matrin 3 is not reported in this study), and then with goat anti-rabbit Dylight 680 and goat anti-moouse Dylight 800. Images on the left represent the rabbit (upper) and mouse (lower) blots from the NONO and SFPQ analysis. Images on the right represent the rabbit (upper) and mouse (lower) blots from the PSPC1 and FUS analysis. The first two lanes in each fraction/lysate are the NTg examples provided in figure 3b.

Supplementary Figure 9

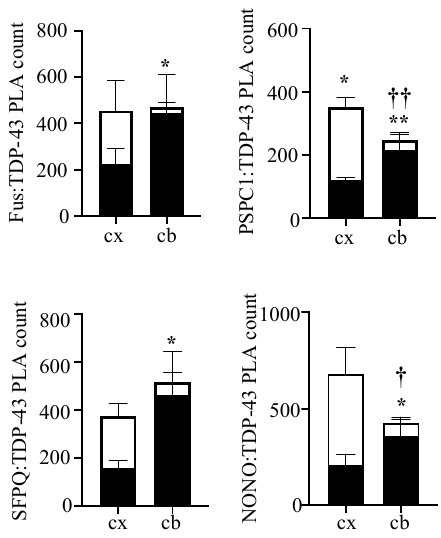

Supplementary Figure 9: Quantification of PLA counts for TDP-43 with FUS, PSPC1, SFPQ and NONO in the nuclear and cytoplasmic regions of the cortex and cerebellum.

There were distinct differences in PLA counts between the granule cell and Purkinje cell layers of the cerebellum, and for logistical reasons these two layers were analysed individually. Since granule cells are the overwhelmingly predominant cells in the cerebellum (~80% of total cell number (Lange, 1975; Nairn et al., 1989)), total PLA count for the cerebellum was calculated as (Purkinje cell layer count x0.2)+(granule cell later count x0.8)

Two-way ANOVA followed by Fisher LSD post hoc test detected a significant statistical interaction between brain region and cellular localisation for PSPC1 (p=0.0018), SFPQ (p=0.0449) and NONO (p=0.0230). No significant interaction was detected with FUS (p=0.0954). Interactions with all paraspeckle proteins were significantly higher in the nuclear fraction than the cytoplasmic fraction in the cerebellum, with the inverse (cytoplasmic interactions higher than nuclear ones) observed for PSPC1 in the cortex. Cytoplasmic interactions with both PSPC1 and NONO were significantly reduced within the cerebellum compared to the cortex.

*p<0.05; **p<0.01 cytoplasmic vs nuclear interactions for the same sample

**†**p<0.05; **††** p<0.01cytoplasmic interactions vs cortex

*Lange, W. (1975) ‘Cell Number and Cell Density in the Cerebellar Cortex of Man and Some Other Mammals’, Cell and Tissue Research, 157(1), pp.115-24. https://doi.org/ 10.1007/BF00223234*

*Nairn, J.G. et al. (1989) ‘On the number of purkinje cells in the human cerebellum: Unbiased estimates obtained by using the “fractionator”’, Journal of Comparative Neurology, 290(4), pp. 527–532. https://doi.org/10.1002/cne.902900407.*
